# Lactate and Immunomagnetic-purified iPSC-derived Cardiomyocytes Generate Comparable Engineered Cardiac Tissue Constructs

**DOI:** 10.1101/2023.05.05.539642

**Authors:** Kalina J. Rossler, Willem J. de Lange, Morgan W. Mann, Timothy J. Aballo, Jake A. Melby, Jianhua Zhang, Gina Kim, Elizabeth F. Bayne, Yanlong Zhu, Emily T. Farrell, Timothy J. Kamp, J. Carter Ralphe, Ying Ge

## Abstract

Three-dimensional engineered cardiac tissue (ECT) using purified human induced pluripotent stem cell–derived cardiomyocytes (hiPSC-CMs) has emerged as an appealing model system for the study of human cardiac biology and disease. A recent study reported widely-used metabolic (lactate) purification of monolayer hiPSC-CM cultures results in an ischemic cardiomyopathy-like phenotype compared to magnetic antibody-based cell sorting (MACS) purification, complicating the interpretation of studies using lactate-purified hiPSC-CMs. Herein, our objective was to determine if use of lactate relative to MACs-purified hiPSC-CMs impacts the properties of resulting hiPSC-ECTs. Therefore, hiPSC-CMs were differentiated and purified using either lactate-based media or MACS. After purification, hiPSC-CMs were combined with hiPSC-cardiac fibroblasts to create 3D hiPSC-ECT constructs maintained in culture for four weeks. There were no structural differences observed, and there was no significant difference in sarcomere length between lactate and MACS hiPSC-ECTs. Assessment of isometric twitch force, Ca^2+^ transients, and β-adrenergic response revealed similar functional performance between purification methods. High-resolution mass spectrometry (MS)-based quantitative proteomics showed no significant difference in any protein pathway expression or myofilament proteoforms. Taken together, this study demonstrates lactate- and MACS-purified hiPSC-CMs generate ECTs with comparable molecular and functional properties, and suggests lactate purification does not result in an irreversible change in hiPSC-CM phenotype.

**Graphical Abstract:** 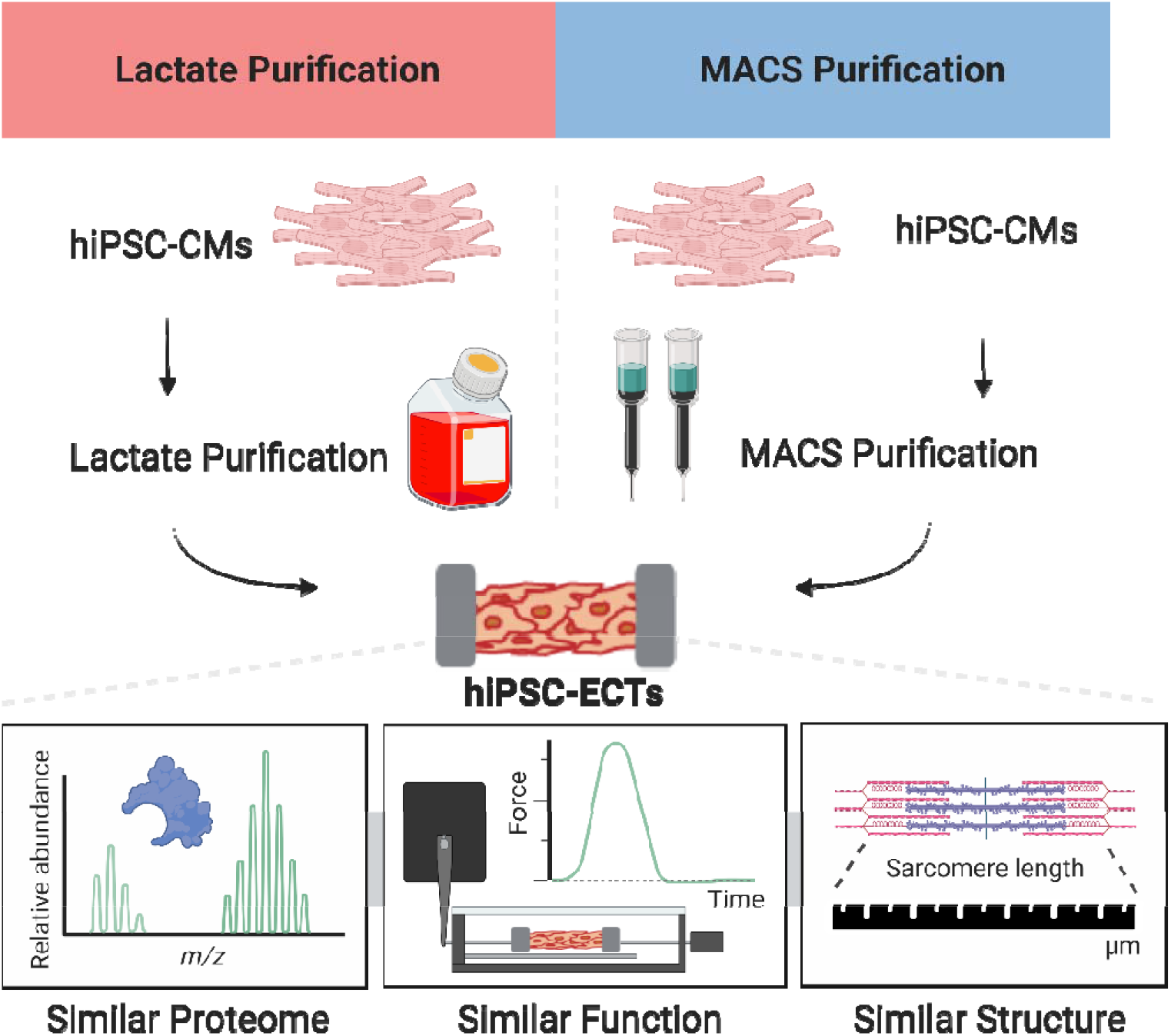

## Introduction

Human induced pluripotent stem cell–derived cardiomyocytes (hiPSC-CMs) have emerged as an important *in vitro* tool for cardiovascular disease modeling, cardiotoxicity screening, and drug discovery.^1,2^ Both monogenic and acquired cardiac diseases have been modeled using two-dimensional (2D) hiPSC-CMs.^3–6^ While hiPSC-CMs permit biological studies in a fully human context, the relative immaturity of hiPSC-CMs remains a notable limitation when considering their application.^7,8^ Significant efforts have been devoted to improving the developmental and maturation state of hiPSC-CM models, including, but not limited to: extending time in culture, treating with specific growth factors, applying mechanical and electrical stimulation, co-culturing with other cell types, and seeding cells in three-dimensional (3D) matrices. ^9–11^ Recent studies showed co-culturing purified hiPSC-CMs with cardiac fibroblasts within three-dimensional engineered cardiac tissue (hiPSC-ECT) constructs improved maturation and function relative to 2D monolayer cultures. The hiPSC-ECT showed evidence of t-tubule systems, more mature intracellular calcium handling, and protein isoform switching – all of which are consistent with more mature heart tissue.^12,13^ Thus, hiPSC-ECTs have great potential for enhanced *in vitro* cardiac modeling.

The first step in hiPSC-ECT generation is the creation of a pure hiPSC-CM population. Purification after hiPSC-CM differentiation is necessary to ensure a more uniform final cell composition within the modeling matrix.^14–16^ This is achieved either by immunomagnetic means – labeling cells using a biotin conjugate and passing through a microbead column (magnetic-activated cell sorting; MACS) – or metabolic purification using a lactate-based media.^17^ To date, the most popular and cost-effective method for the purification of 2D hiPSC-CMs is the use of glucose-free, lactate media. This approach relies on the unique ability of CMs to effectively utilize lactate as a metabolic substrate over other cells.^18^ Thus, lactate purification is currently preferred by many laboratories for generation of 2D and 3D hiPSC-CM models.^15,19–39^ Notably, lactate-purified hiPSC-CMs have been used to test new therapeutic agents for diseases such as Duchenne muscular dystrophy^34^ and Lamin induced cardiomyopathies.^40^

Despite its widespread use, questions regarding the metabolic stress caused by lactate selection have been raised. A recent study by Davis et al. suggests lactate purification of 2D hiPSC-CM cultures results in an “ischemic-like” phenotype modeling ischemic heart failure *in vitro*, making them less desirable for widespread applications.^3^ The lactate-purified hiPSC-CMs exhibited changes in molecular composition, structure, and function relative to MACS purified hiPSC-CMs. For example, lactate-purified hiPSC-CMs displayed spontaneous arrhythmic activity, abnormal calcium handling, shortened sarcomeres, and aberrant contractile protein expression. Based on the concerns raised by this work, our objective was to investigate if the responses to lactate purification identified in Davis et. al in 2D-cultured hiPSC-CMs represented a transient metabolic adaptation or a persistent phenotypic change. We hypothesized prolonged culture of lactate- or MACS-purified hiPSC-CMs in a 3D environment in ECTs for four weeks would generate in comparable functioning cardiac tissue.

To investigate our hypothesis, we compared the structure, function, and protein expression of hiPSC-ECTs containing either lactate-purified hiPSC-CMs or hiPSC-CMs purified using a commercially available MACS assay (Miltenyi Biotec). We observed no differences in overall structure and performed immunohistochemical staining of the cardiac Z-discs to quantify sarcomere length. We assessed the functional performance of our hiPSC-ECTs in both the absence and presence of beta-adrenergic stimulation. Moreover, we utilized an unbiased high-resolution mass-spectrometry-based proteomics method that integrated both bottom-up global proteomics to comprehensively assess global proteome changes and top-down targeted proteomics to accurately determining the changes in sarcomeric post-translational modifications (PTMs) and isoforms, collectively called proteoforms.^13,41^ This detailed analysis of structure, function, and proteome revealed no significant differences between lactate and MACS hiPSC-ECTs. Taken together, this study suggests the method of purification of cardiomyocytes prior to utilizing hiPSC-CMs in 3D culture platforms does not significantly influence important functional and molecular characteristics.

## Results

### Generation of hiPSC-ECT constructs from purified hiPSC-CMs

We utilized an established control lineage (Stanford University Cardiovascular Institutes Biobank; Hypertrophic Cardiomyopathy/*MYH7*/Control iPSC line) and a monolayer-based, bi-phasic modulation of Wnt signaling protocol (GiWi protocol)^4,42^ to generate hiPSC-CMs for our study (Figure 1A). We then enriched our differentiated hiPSC-CM cultures using either a lactate-based media or magnetic antibody-based (MACS) depletion of the non-myocyte cell populations. Each batch of hiPSC-CMs was split with cells from each differentiation applied to both arms (lactate vs. MACS) of our study to ensure well-matched comparisons. After purification, we combined hiPSC-CMs in a 10:1 ratio with low passage isogenic hiPSC-cardiac fibroblasts (CFs) and seeded the cell mixture in a fibrinogen-thrombin matrix to form hiPSC-ECTs.^43^ We then cultured the hiPSC-ECTs for four weeks to allow for construct remodeling and maturation before subsequent analyses. Notably, hiPSC-CM purification using both methods yielded similar percentages of CMs (Figure 1B) as previously described.^3^ Recovery of pure hiPSC-CMs was greater using lactate purification versus MACS (Supplemental Figure 1; lactate = 82.21 ± 2.86 %, n = 4 vs. MACS = 60.87 ± 8.73 %, n = 4; p = 0.0035). No visual differences were observed between lactate and MACS hiPSC-ECTs, indicating successful remodeling in both constructs (Figure 1C).

**Figure 1.**
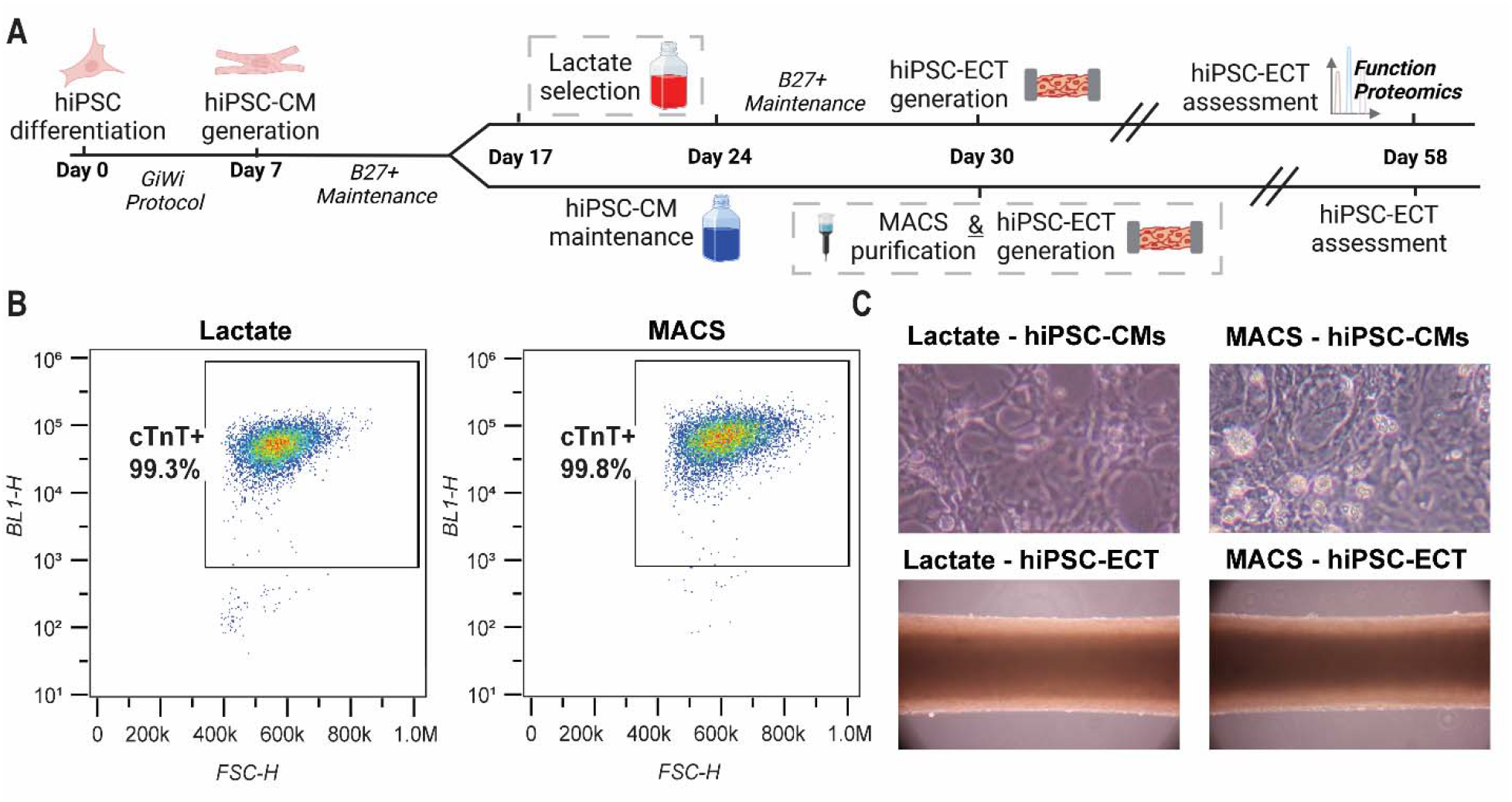
Purification of 2D hiPSC-CMs and 3D hiPSC-ECT generation. (A) Timeline of 3D human induced pluripotent stem cell (hiPSC)-derived engineered cardiac tissues (ECTs). (B) Flow cytometry with cTnT labeling demonstrates effective hiPSC-CM enrichment using each purification method. (C) 2D and 3D representations of hiPSC-CMs from lactate and MACS purification methods (10X objective).

### hiPSC-ECT structure is consistent between lactate and MACS purification methods

After seeing no differences between hiPSC-ECTs using a microscope, we assessed the myofibril structure of our hiPSC-ECTs using histological methods. After 4 weeks in culture, a separate cohort of hiPSC-ECTs were fixed in OCT, sectioned, and stained with alpha-actinin, a Z-disc protein flanking the ends of the cardiac sarcomere, to assess sarcomere length (Figure 2A; Supplemental Figure 2).^44,45^ Three hiPSC-ECTs per condition was sectioned into multiple slides to assess sarcomere structure and cellular alignment per purification method. Indeed, there was no statistical difference in comparison to lactate hiPSC-ECTs (Figure 2B; lactate = 1.67 ± 0.02 µm, n = 99 vs. MACS = 1.65 ± 0.02 µm, n = 100; unpaired t test, p = 0.4460). This analysis taken together indicates 2D hiPSC-CM purification methods do not affect myofibril organization in 3D hiPSC-ECTs.

**Figure 2.**
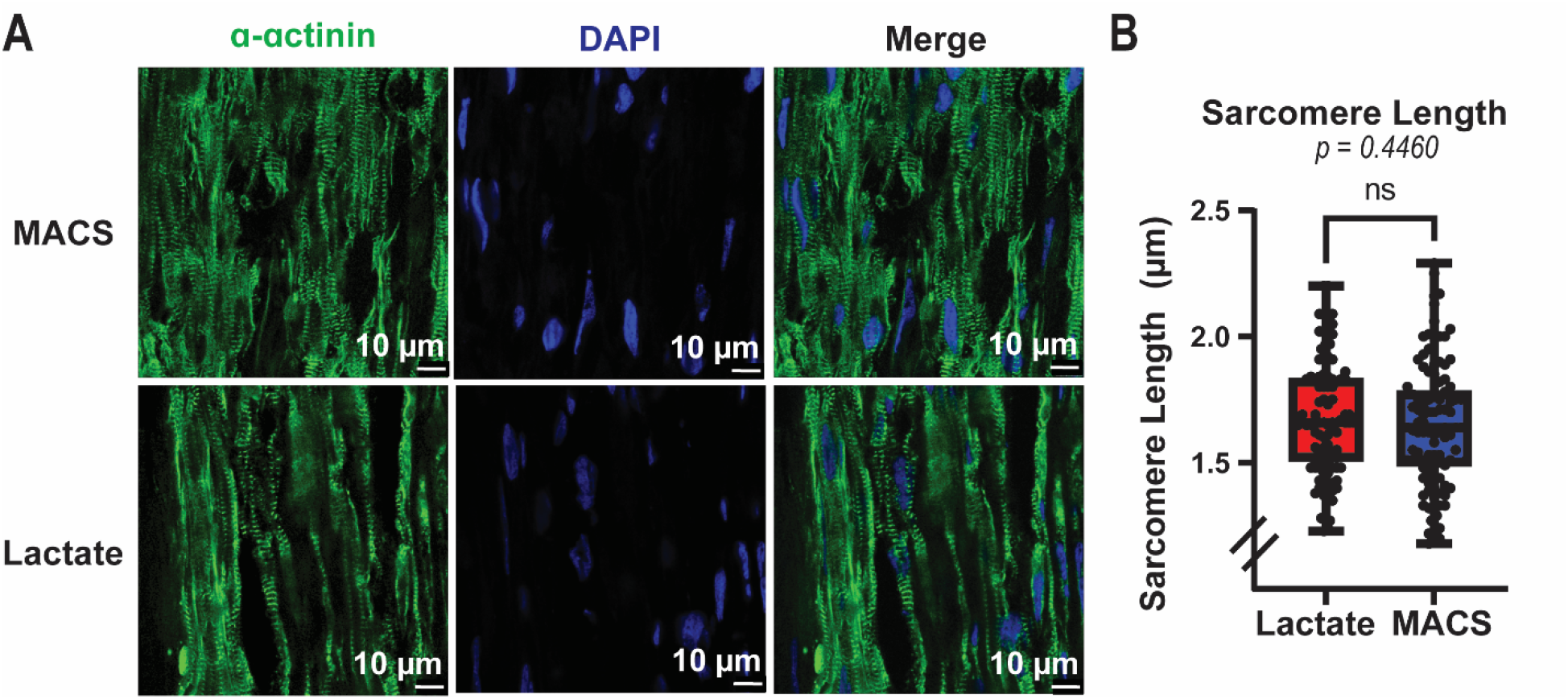
Sarcomere lengths of hiPSC-ECTs. (A) Alpha-actinin and DAPI immunofluorescent labeling on representative lactate and MACS hiPSC-ECTS. (B) Sarcomere length comparison (lactate = 1.67 ± 0.02 µm, n = 99 vs. MACS = 1.65 ± 0.02 µm, n = 100; unpaired t test, p = 0.4460). All statistical analyses are unpaired t tests with α = 0.05. Plots show whiskers from 0-25^th^ percentile, box from 25-75^th^ percentile (mean indicated with line), and whiskers from 75-100^th^ percentile.

### Lactate purification has no impact on hiPSC-ECT twitch force parameters compared to MACS hiPSC-ECTs

After 4 weeks in 3D culture, we performed extensive functional characterization of the lactate and MACS hiPSC-ECTs. hiPSC-ECT constructs were electrically paced at 1.5 Hz in a 37 °C physiologic chamber for the duration of the functional analyses. To determine baseline contractile function, we measured twitch force amplitude (TF), time from pacing stimulus to twitch force peak (CT_100_), time from twitch force peak to 50% twitch force decay (RT_50_), and time from 50% to 90% twitch force decay (RT_50–90_) (Figure 3; Supplemental Figure 3A). Cross sectional area (CSA) was not significantly different (Figure 3C; lactate = 0.11 ± 0.06 mm^2^, n = 7 vs. MACS = 0.15 ± 0.04 mm^2^, n = 5; unpaired t test, p = 0.2536) indicating successful construct remodeling in both MACS and lactate hiPSC-ECTs.

**Figure 3.**
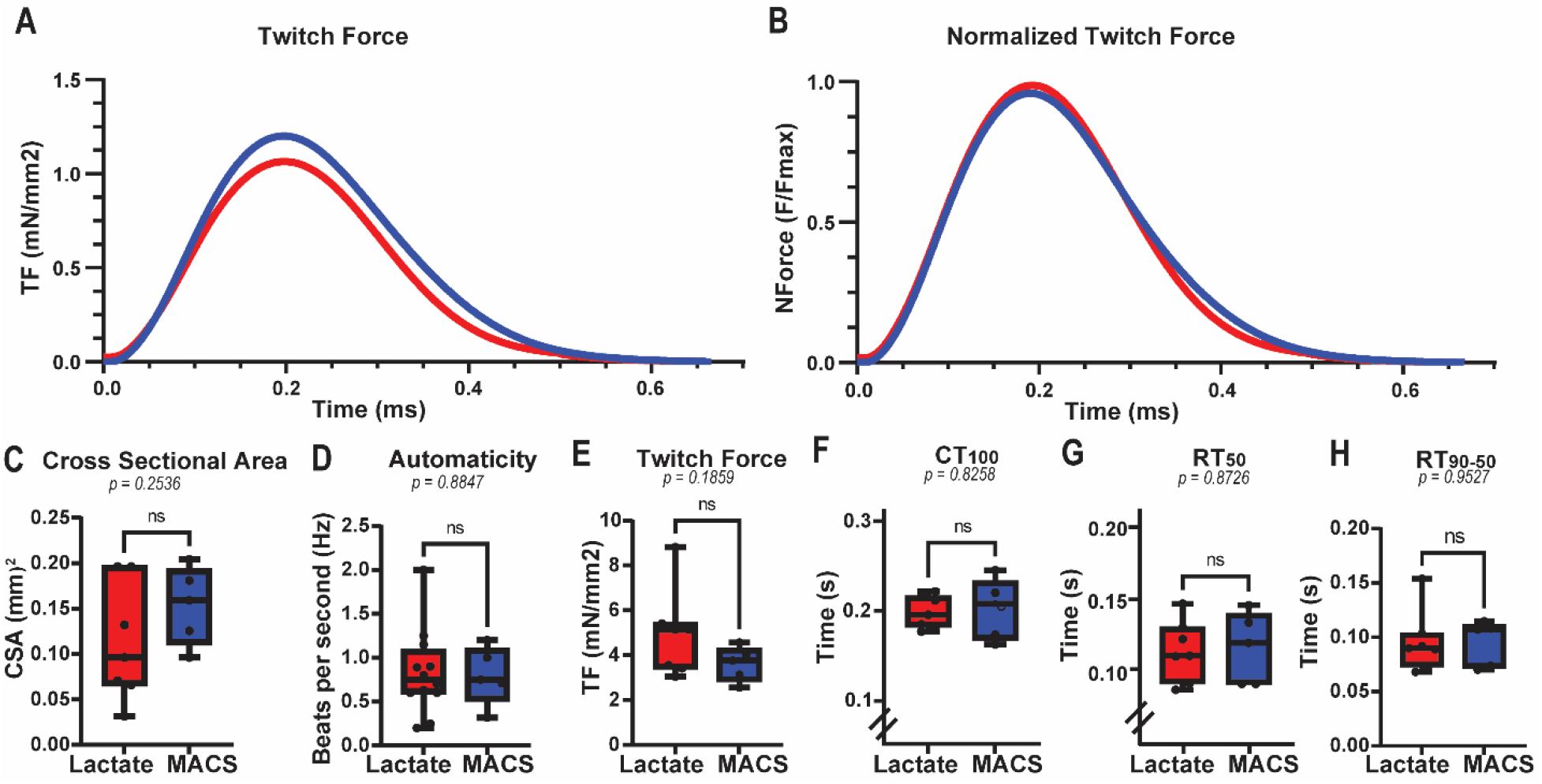
Twitch Force Assessment of hiPSC-ECTs. (A) Raw and (B) normalized averaged twitch force curves for lactate (red) and MACS (blue) hiPSC-ECTs. (C-H) Twitch force parameters for lactate (red) and MACS (blue) hiPSC-ECTs as follows: (C) cross sectional area (CSA; lactate = 0.11 ± 0.06 mm^2^, vs. MACS = 0.15 ± 0.04 mm^2^, p = 0.2536); (D) automaticity (lactate = 0.83 ± 0.14 Hz, n = 12 vs. MACS = 0.80 ± 0.15 Hz, n = 5; p = 0.8847); (E) twitch force amplitude (TF; lactate = 4.93 ± 0.74 mN/mm^2^, vs. MACS = 3.60 ± 0.35 mN/mm^2^, p = 0.1859); (F) time from pacing stimulus to twitch force peak (CT_100_; lactate = 0.200 ± 0.01 s, vs. MACS = 0.20 ± 0.02 s, p = 0.8258); (G) time from twitch force peak to 50% twitch force decay (RT_50_; lactate = 0.11 ± 0.01 s, vs. MACS = 0.12 ± 0.01 s, p = 0.8726); and (H) time from 50% to 90% twitch force decay (RT_50–90_; lactate = 0.10 ± 0.01 s, vs. MACS = 0.09 ± 0.09 s, p = 0.9527). All tests performed with biological replicates as lactate, n=7 and MACS, n=5, unless otherwise stated. All statistical analyses are unpaired t tests with α = 0.05. Plots show whiskers from 0-25^th^ percentile, box from 25-75^th^ percentile (mean indicated with line), and whiskers from 75-100^th^ percentile.

Additionally, automaticity was not significantly different between enrichment conditions (Figure 3D; lactate = 0.83 ± 0.14 Hz, n = 12 vs. MACS = 0.80 ± 0.15 Hz, n = 5; unpaired t test, p = 0.8847), and no arrythmias were observed while capturing automaticity (Supplemental Figure 4). Twitch force amplitude was similar for lactate and MACS hiPSC-ECTs (Figure 3E; lactate = 4.93 ± 0.74 mN/mm^2^, n = 7 vs. MACS = 3.60 ± 0.34 mN/mm^2^, n = 5; unpaired t test, p = 0.1859). Other twitch force parameters, such as CT_100_ (Figure 3F; lactate = 0.20 ± 0.01 s, n = 7 vs. MACS = 0.20 ± 0.02 ds, n = 5; unpaired t test, p = 0.8258), RT_50_ (Figure 3G; lactate = 0.11 ± 0.01 s, n = 7 vs. MACS = 0.12 ± 0.01 s, n = 5; unpaired t test, p = 0.8726), and RT_50-90_ (Figure 3H; lactate = 0.10 ± 0.01 s, n = 7 vs. MACS = 0.09 ± 0.09 s, n = 5; unpaired t test, p = 0.9527), displayed similar ranges across hiPSC-ECTs and were not significantly different. Collectively, the contractile measurements indicate lactate hiPSC-ECTs display similar baseline functional properties compared to MACs hiPSC-ECTs.

### Intracellular calcium handling and β-adrenergic response is similar between purification protocols

We then sought to understand how calcium handling may be affected in our lactate hiPSC-ECTs. Calcium transients (Ca^2+^TR) were assessed at 1.5 Hz using the Ca^2+^-indicator Fura2-AM (Figure 4). All Ca^2+^ parameters were consistently similar between lactate and MACS hiPSC-ECTs, including Ca^2+^TR peak (Figure 4B; dR; lactate = 0.41 ± 0.09 Fura340/380, n = 7 vs. MACS = 0.67 ± 0.11 Fura340/380, n = 5; unpaired t test, p = 0.0962), time to Ca^2+^TR peak (Figure 4C; CaT_100_; lactate = 0.11 ± 0.01 s, n = 7 vs. MACS = 0.11 ± 0.01 s, n = 5; unpaired t test, p = 0.8327), time from Ca^2+^TR peak to 50% Ca^2+^TR decay (Figure 4D; CaDT_50_; lactate = 0.19 ± 0.02 s, n = 7 vs. MACS = 0.18 ± 0.01 s, n = 5; unpaired t test, p = 0.5104), and time from 50% Ca^2+^TR decay to 75% Ca^2+^TR decay (Figure 4E; CaDT_50-75_; lactate = 0.11 ± 0.01 s, n = 7 vs. MACS = 0.09 ± 0.01 s, n = 5; unpaired t test, p = 0.0581). These findings indicate Ca^2+^-handling is similar between lactate and MACS hiPSC-ECTs.

**Figure 4.**
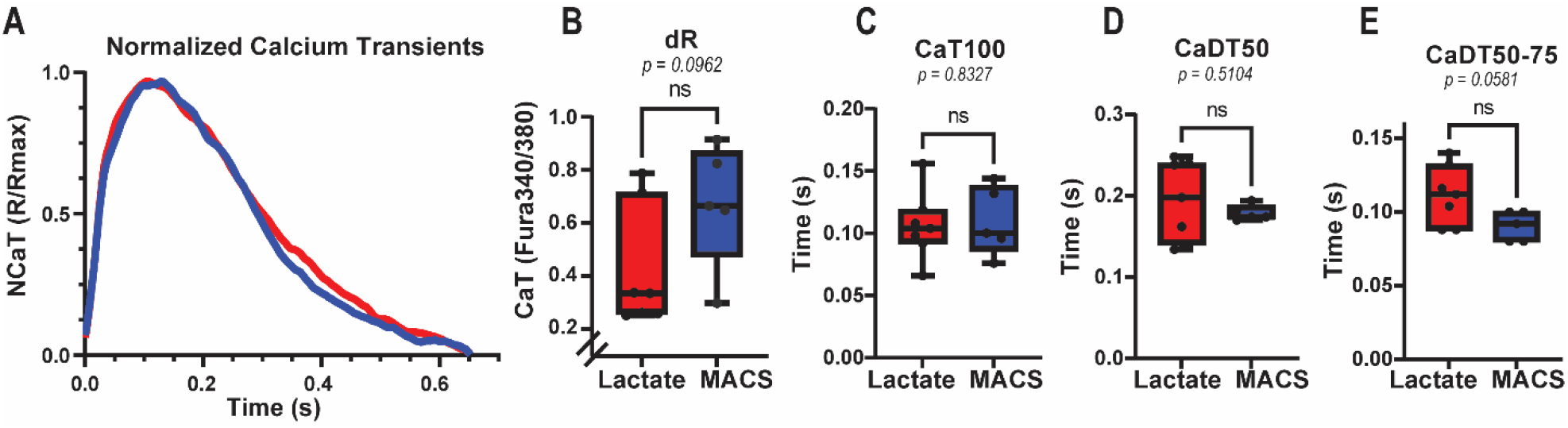
Calcium Transients Assessment of hiPSC-ECTs. (A) Normalized averaged calcium transients (Ca^2+^TR) curves for lactate (red) and MACS (blue) hiPSC-ECTs. (B) Ca^2+^TR parameters for lactate (red) and MACS (blue) hiPSC-ECTs as follows: (B) Ca^2+^TR peak (dR; lactate = 0.41 ± 0.09 Fura340/380, n = 7 vs. MACS = 0.67 ± 0.11 Fura340/380, n = 5; unpaired t test, p = 0.0962); (C) time to Ca^2+^TR peak (CaT_100_; lactate = 0.11 ± 0.01 s, n = 7 vs. MACS = 0.11 ± 0.01 s, n = 5; unpaired t test, p = 0.8327); (D) time from Ca^2+^TR peak to 50% Ca^2+^TR decay (CaDT_50_; lactate = 0.19 ± 0.02 s, n = 7 vs. MACS = 0.18 ± 0.01 s, n = 5; unpaired t test, p = 0.5104); (D) and time from 50% Ca^2+^TR decay to 75% Ca^2+^TR decay (CaDT_50-75_; lactate = 0.11 ± 0.01 s, n = 7 vs. MACS = 0.09 ± 0.01 s, n = 5; unpaired t test, p = 0.0581). All tests performed with biological replicates as lactate, n=7 and MACS, n=5, unless otherwise states. All statistical analyses are unpaired t tests with α = 0.05. Plots show whickers from 0-25^th^ percentile, box from 25-75^th^ percentile (mean indicated with line), and whiskers from 75-100^th^ percentile.

To probe beta-adrenergic responsiveness of our hiPSC-ECTs, we treated our constructs with 1□µM isoproterenol for 5□min (Supplemental Figure 5). Isoproterenol treatment positively increased the automaticity of all hiPSC-ECTs. The percentage increase given by beta-adrenergic response was similar between lactate and MACS hiPSC-ECTs (lactate = 27.28 ± 5.87 %, n=8 vs. MACS = 13.35 ± 2.24 %, n=5; unpaired t test, p = 0.1965), indicating a positive chronotropic effect in both groups. Notably, no arrythmias were observed while capturing automaticity post-treatment (Supplemental Figure 4). There was no significant difference between lactate and MACS hiPSC-ECTs in the percentage given by isoproterenol effect on either twitch force parameters (TF; lactate = 3.85 ± 3.44 %, n = 7 vs. MACS = 8.35 ± 2.43 %, n = 5; unpaired t test, p = 0.3508; CT_100_; lactate = -2.14 ± 1.79 %, n = 7 vs. MACS = -2.18 ± 1.69 %, n = 5; unpaired t test, p = 0.9862; RT_50_; lactate = -9.70 ± 3.14 %, n = 7 vs. MACS = -6.24 ± 3.37 %, n = 5; unpaired t test, p = 0.4776; RT_50-90_; lactate = -7.12 ± 5.26 %, n = 7 vs. MACS = -2.22 ± 3.63 %, n = 5; unpaired t test, p = 0.5061) or calcium transients assessments (dR; lactate = 27.43 ± 6.07 %, n = 7 vs. MACS = 64.84 ± 42.92 %, n = 5; unpaired t test, p = 0.3267; CaT_100_; lactate = -9.41 ± 5.30 %, n = 7 vs. MACS = -5.24 ± 5.99 %, n = 5; unpaired t test, p = 0.6163; CaDT_50_; lactate = -3.54 ± 5.26 %, n = 7 vs. MACS = -20.47 ± 9.14 %, n = 5; unpaired t test, p = 0.1164; CaDT_50-75_; lactate = -6.78 ± 6.15 %, n = 7 vs. MACS = 15.66 ± 14.90 %, n = 5; unpaired t test, p = 0.1496). Taken together, these data suggest there is similar function between lactate and MACS hiPSC-ECTs.

### Global proteome of lactate hiPSC-ECTs is similar to MACS hiPSC-ECTs

After functional characterization, we sought to assess the global expression profile arising from the same hiPSC-ECTs. Immediately following isoproterenol treatment, hiPSC-ECTs were flash frozen and stored at -80°C until proteomic analysis. A buffer containing our in-house photocleavable surfactant, Azo (0.25% w/v), was used to extract the global proteome similar to established protocols.^46,47^ Aliquots from the protein homogenate were saved for SDS-page analysis of extraction reproducibility (Supplemental Figure 6), and instrument reproducibility was evaluated using total ion chromatograms (Supplemental Figure 7).

We identified over 5000 unique proteins in each hiPSC-ECT sample (Figure 5A; lactate = 5299 ± 60 proteins, n = 7 vs. MACS = 5262 ± 28 proteins, n = 5). Pearson correlations of overall protein expression revealed the greatest deviation from linearity between any pair of hiPSC-ECT as R = 0.94. Unbiased hierarchical clustering of hiPSC-ECT over all protein expression indicated via dendrogram randomly grouped replicates between lactate and MACS hiPSC-ECTs (Figure 5B). Visualization of overall protein group expression between purification techniques did not reveal any differentially expressed proteins (Figure 5C).

**Figure 5.**
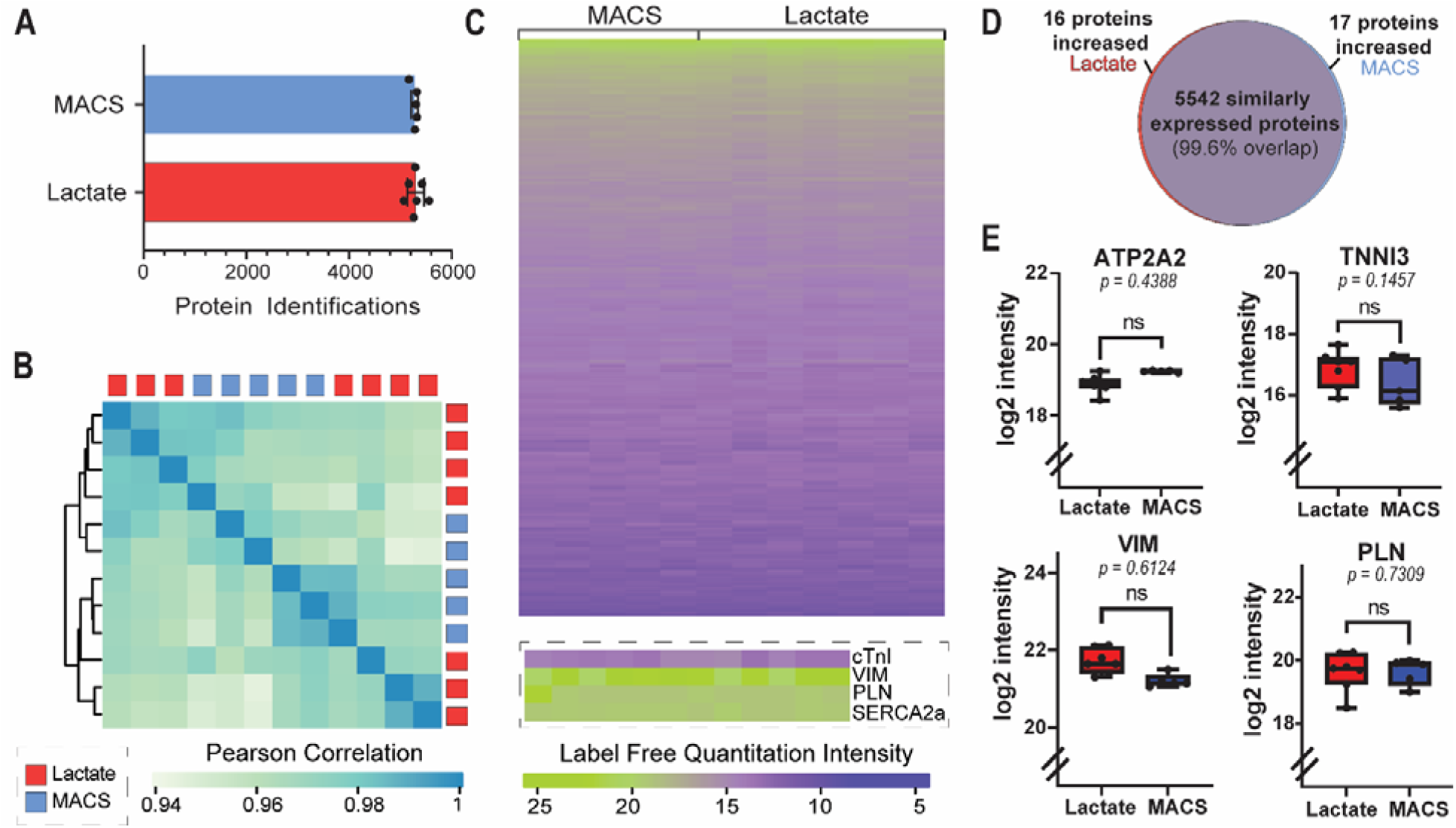
Global proteomics analysis of hiPSC-ECTs. (A) Unique protein identifications per hiPSC-ECT group (lactate = 5299 ± 61 proteins, n = 7 vs. MACS = 5262 ± 28 proteins, n = 5). (B) Pearson correlation of hiPSC-ECT replicates with unbiased dendrogram clustering. (C) Visual representation of differentially expressed proteins. (D) Heat map of overall protein expression with expression of cardiac troponin I (cTnI), phospholamban (PLN), cardiac sarcoplasmic reticulum Ca^2+-^ATPase2a (SERCA2a), and vimentin (VIM) shown for each replicate. (E) Log_2_ fold intensity values plotted for cTnI, PLN, SERCA2a, and VIM, as proteins of interest (cTnI; lactate = 16.86 ± 0.22 log_2_ fold value, n = 7 vs. MACS = 16.41 ± 0.34 log_2_ fold value, n = 5; unpaired t test, p = 0.1457; PLN; lactate = 19.64 ± 0.23 log_2_ fold value, n = 7 vs. MACS = 19.64 ± 0.19 log_2_ fold value, n = 5; unpaired t test, p = 0.7309; SERCA2a; lactate = 18.91 ± 0.10 log_2_ fold value, n = 7 vs. MACS = 19.26 ± 0.01 log_2_ fold value, n = 5; unpaired t test, p = 0.4388; VIM; lactate = 21.72 ± 0.12 log_2_ fold value, n = 7 vs. MACS = 21.21 ± 0.08 log_2_ fold value, n = 5; unpaired t test, p = 0.6124). Plots show whiskers from 0-25^th^ percentile, box from 25-75^th^ percentile (mean indicated with line), and whiskers from 75-100^th^ percentile.

Next, we performed differential expression analysis to identify proteins and/or pathways that were changing between hiPSC-ECT groups (adjusted p-value of 0.05 or less and a log_2_ fold change of 1 for each protein). Our differential expression protein analysis revealed 5542 proteins commonly expressed in all replicates with only 33 proteins being differentially expressed (99.6% similarity; Figure 5D, Supplemental Figure 8).

We performed pathway analyses using KEGG, UniProt, and PANTHER databases to identify cellular and disease pathway trends within differentially expressed proteins.^48–50^ None of the databases indicated significantly changing pathways for the differentially expressed proteins. Therefore, we manually searched for overall pathway protein expression differences in our dataset using relevant keywords such as “muscle”, “cardiomyopathy”, “reactive oxygen species”, “glycolysis”, “beta-oxidation”, “microtubule”, “hypoxia”, “senescence”, “TGFβ”, “adrenaline”, “trafficking”, “hypertrophy,” and “mitochondrial”. These pathways were plotted with all identified proteins (Supplemental Figure 9). Overall, there was no significant changes in protein expression for the pathways identified. Key proteins identified by Davis et al. as ischemic markers, including cardiac troponin I (cTnI), phospholamban (PLN), cardiac sarcoplasmic reticulum Ca^2+-^ATPase2a (SERCA2a), and vimentin (VIM), were not differentially expressed (Figure 5E; cTnI; lactate = 16.86 ± 0.23 log_2_ fold value, n = 7 vs. MACS = 16.41 ± 0.34 log_2_ fold value, n = 5; unpaired t test, p = 0.1457; PLN; lactate = 19.64 ± 0.23 log_2_ fold value, n = 7 vs. MACS = 19.64 ± 0.19 log_2_ fold value, n = 5; unpaired t test, p = 0.7309; SERCA2a; lactate = 18.91 ± 0.10 log_2_ fold value, n = 7 vs. MACS = 19.26 ± 0.01 log_2_ fold value, n = 5; unpaired t test, p = 0.4388; VIM; lactate = 21.72 ± 0.12 log_2_ fold value, n = 7 vs. MACS = 21.21 ± 0.08 log_2_ fold value, n = 5; unpaired t test, p = 0.6124).^3^ Therefore, our proteomics data indicate global proteome expression is unaffected by lactate purification.

### Myofilament proteoform expression of lactate hiPSC-ECTs is comparable to MACS hiPSC-ECTs

We then sought to evaluate the possible effect of lactate purification on myofibril organization through sarcomere protein expression and organization. We aimed to target possible protein isoform and phosphorylation changes within the proteins of the contractile apparatus using intact protein analysis. After baseline twitch force characterization only, a separate group of hiPSC-ECTs was flash-frozen and stored at −80 °C until targeted top-down proteomics analysis was performed as previously described.^51^ For myofilament analysis, constructs were homogenized in a lysis buffer containing HEPES (pH = 7) to deplete cytosolic proteins. The sarcomere-enriched pellet was exposed to a trifluoroacetic acid (TFA) buffer (pH = 2) to create a sarcomere-rich extract. Equal amounts of proteins from each extract were analyzed by top-down LC-MS/MS (Supplemental Figure 10).

Our highly reproducible, top-down targeted proteomics allowed us to obtain a bird’s eye view of all the detectable PTMs and isoforms of sarcomere proteins in a single injection (Figure 6A; Supplemental Figure 11-13).^13,51–53^ Due to the ability of intact protein analysis to confidently differentiate between isoforms, we quantified several sarcomere isoforms to validate our global proteomics results. Myosin light chain-2v (MLC-2v) is a crucial component of the myosin motor and is directly involved in sarcomere contraction.^54^ Its emerging expression is indicative of a “mature” ventricular-like phenotype in hiPSC-CMs.^13,42^ As anticipated, the expression of MLC-2v given by the area under the curve (AUC) was comparable between lactate and MACS hiPSC-ECTs (Figure 6B; lactate = 2.46 x 10^7^ ± 6.27 x 10^6^ AUC, n = 6 vs. MACS = 1.01 x 10^7^ ± 1.19 x 10^6^ AUC, n=3; unpaired t test, p = 0.1604). Furthermore, the phosphorylation of MLC-2v was found to be similar by pTotal calculations (*pTotal* = mol of phosphorylated isoform/mol of total protein; lactate = 0.018 ± 0.00 pTotal, n = 6 vs. MACS = 0.023 ± 0.01 pTotal, n = 3; unpaired t test, p = 0.5008). Unphosphorylated and phosphorylated states of contractile thin filament protein alpha-tropomyosin (α-Tpm) were identified in both lactate and MACS hiPSC-ECTs. A-Tpm was the only tropomyosin isoform detected, and the total expression and phosphorylation of α-Tpm was not significantly different (Figure 6C; lactate = 1.12 x 10^6^ ± 6.30 x 10^4^ AUC, n = 6 vs. MACS = 9.05 x 10^5^ ± 1.29 x 10^5^ AUC, n=3; unpaired t test, p = 0.1365; lactate = 0.619 ± 0.05 pTotal, n = 6 vs. MACS = 0.670 ± 0.03 pTotal, n = 3; unpaired t test, p = 0.5237). Slow skeletal troponin I (ssTnI) is a critical isoform in cardiac development and is often used in ratio to cardiac troponin I (cTnI) to evaluate model maturation.^13,55^ While cTnI has been previously reported using our top-down method for hiPSC-ECTs, its expression was not detected with this cell line.^51^ Nevertheless, the relative expression of ssTnI did not change between hiPSC-ECT groups (Figure 6D; lactate = 3.56 x 10^6^ ± 5.94 x 10^5^ AUC, n = 6 vs. MACS = 2.61 x 10^5^ ± 2.62 x 10^5^ AUC, n = 3; unpaired t-test, p = 0.2961). Differential expression of myosin light chain (MLC) isoform can often indicate differences in maturation state.^13^ We did not identify significant changes in any myosin light chain isoforms, including MLC-1a (Figure 6E; lactate = 1.96 x 10^7^ ± 4.20 x 10^6^ AUC, n = 6 vs. MACS = 1.11 x 10^7^ ± 1.01 x 10^6^ AUC, n = 3; unpaired t-test, p = 0.2108), MLC-1v (lactate = 7.57 x 10^6^ ± 1.46 x 10^6^ AUC, n = 6 vs. MACS = 2.77 x 10^6^ ± 8.51 x 10^5^ AUC, n = 3; unpaired t-test, p = 0.0667), or their isoform ratio (lactate = 2.70 ± 0.310 AUC MLC-1a/1-v, n = 6 vs. MACS = 4.94 ± 1.54 AUC MLC-1a/1-v, n = 3; unpaired t-test, p = 0.0827). Cardiac troponin T (cTnT6) modulates the interaction between the thick and thin filaments in the presence of calcium, and likewise, the phosphorylation state was not statistically different (Supplemental Figure 13; lactate = 0.616 ± 0.05 pTotal, n = 6 vs. MACS = 0.598 ± 0.15 pTotal, n = 3; unpaired t test, p = 0.8878).^56^ The absences of a significant functional differences between ECT formed from either lactate- or MACS-purified hiPSC-CMs is supported at the level of the proteome.

**Figure 6.**
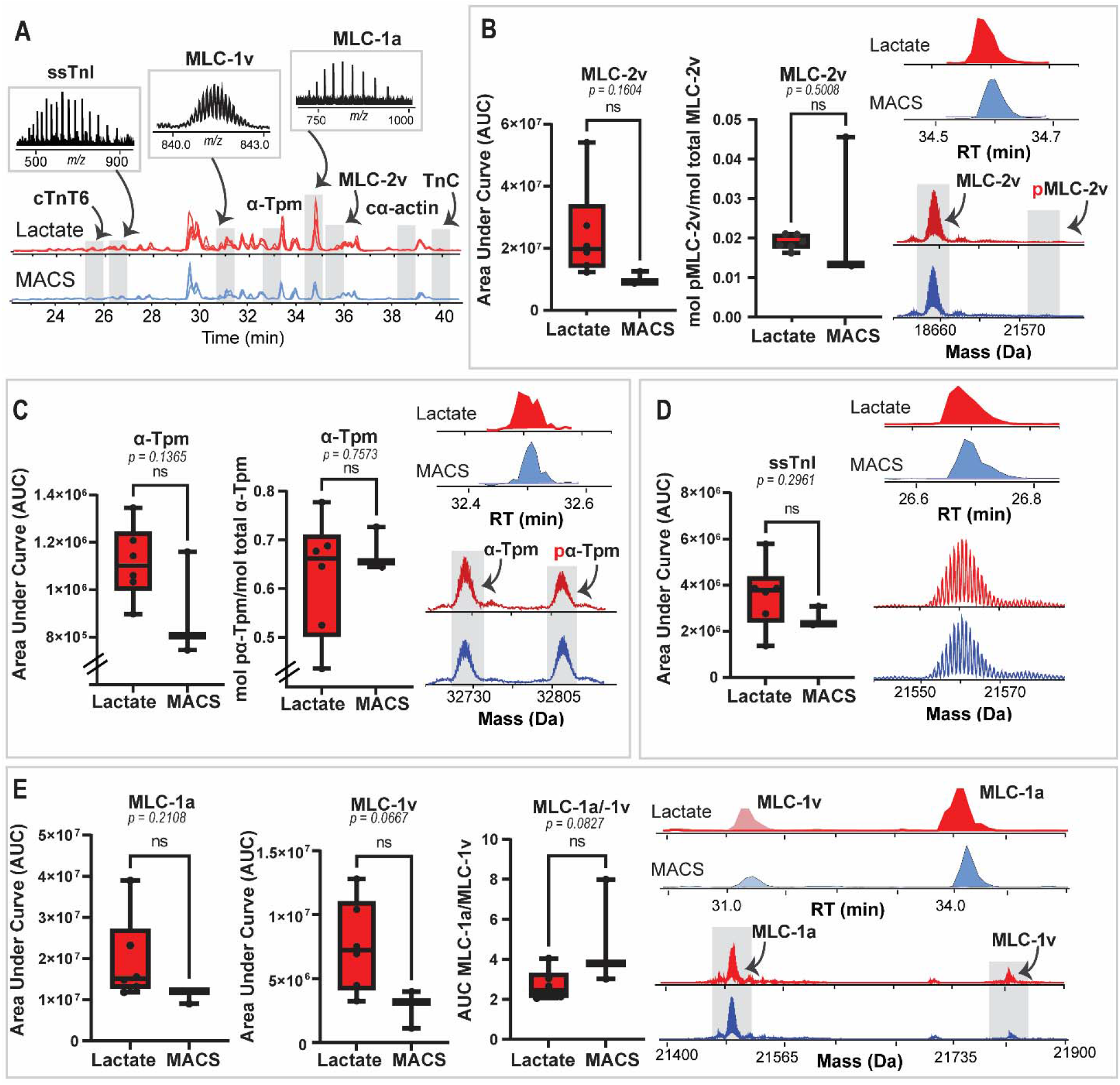
Top-down proteomics of the cardiac sarcomere. (A) Selected base peak chromatograms for lactate and MACS replicates with identified proteins based on retention time. (B) Myosin light chain-2v (MLC-2v) quantitation and phosphorylation (lactate = 2.46 x 10^7^ ± 6.27 x 10^6^ AUC, n = 6 vs. MACS = 1.02 x 10^7^ ± 1.19 x 10^6^ AUC, n=3; unpaired t test, p = 0.1604; lactate = 0.018 ± 0.00 pTotal, n = 6 vs. MACS = 0.023 ± 0.01 pTotal, n = 3; unpaired t test, p = 0.5008). (C) Alpha-tropomyosin (α-Tpm) quantitation and phosphorylation (lactate = 1.12 x 10^6^ ± 6.30 x 10^4^ AUC, n = 6 vs. MACS = 9.05 x 10^5^ ± 1.29 x 10^5^ AUC, n=3; unpaired t test, p = 0.137; lactate = 0.619 ± 0.05 pTotal, n = 6 vs. MACS = 0.670 ± 0.03 pTotal, n = 3; unpaired t test, p = 0.5237).b(D) Slow skeletal troponin I (ssTnI) quantitation (lactate = 3.56 x 10^6^ ± 5.94 x 10^5^ AUC, n = 6 vs. MACS = 2.61 x 10^5^ ± 2.62 x 10^5^ AUC, n = 3; unpaired t-test, p = 0.2961). (E) Myosin light chain isoform quantitation. (MLC-1;lactate = 1.96 x 10^7^ ± 4.20 x 10^6^ AUC, n = 6 vs. MACS = 1.11 x 10^7^ ± 1.01 x 10^6^ AUC, n = 3; unpaired t-test, p = 0.2108; MLC-1v; lactate = 7.57 x 10^6^ ± 1.46 x 10^6^ AUC, n = 6 vs. MACS = 2.77 x 10^6^ ± 8.51 x 10^5^ AUC, n = 3; unpaired t-test, p = 0.0667; MLC-1a/-1v isoform ratio; lactate = 2.70 ± 0.310 AUC MLC-1a/1-v, n = 6 vs. MACS = 4.94 ± 1.54 AUC MLC-1a/1-v, n = 3; unpaired t-test, p = 0.0827). Ptotal given by mol phosphorylated protein/mol total protein. Retention times shown by extracted ion chromatograms (using top 5 charge states). Deconvoluted spectra shown for quantitated proteins. All statistics given by unpaired t-test with alpha = 0.05. Plots show 0-25^th^ percentile (bottom “whisker”), 25^th^ to 75^th^ percentile (box) with mean indicated using line, and 75^th^ percentile to 100^th^ percentile (top whisker).

## Discussion

Lactate purification of hiPSC-CMs for subsequent use in 2D and 3D modeling has been widely adopted by the cardiovascular research community.^18^ Davis et al. recently provided data suggesting the lactate purification of 2D hiPSC-CMs generates a phenotypically “ischemic” cell population, implying an adverse effect of metabolic selection on cells and therefore, negative implications for their use in certain downstream applications. This is a critical issue since the purified hiPSC-CMs are being utilized by many groups to explore often subtle biological questions.^19–39^ Given the proven benefits of growing hiPSC-CMs in an integrated 3D tissue environment, we explored whether the 3D ECT would mitigate the functional issues observed in 2D-cultured lactate-purified hiPS-CMs compared to the MACS-purification approach. Our complete structural and functional comparison of 3D hiPSC-ECTs generated using either lactate or MACS purified hiPSC-CMs demonstrated a high degree of similarity between groups. Both hiPSC-ECT cohorts were structurally similar under a microscope and at the myofilament level. Functionally, lactate hiPSC-ECTs demonstrated similar contractile properties and calcium handling to our MACS hiPSC-ECTs. Importantly, this phenotype did not change with isoproterenol treatment, with both hiPSC-ECT cohorts displaying a positive chronotropic effect. Changes across the proteome as well as alterations in myofilament proteoforms were not detected.

Remodeling of the 3D hiPSC-ECT construct is a critical component to successful model function and can be determined visually using a microscope.^12,57^ We did not observe any differences in structure between the hiPSC-ECT groups at the macrolevel. A decrease in sarcomere length has been observed previously in animal models with myocardial infarction, specifically within the ischemic regions of the heart^58^. Indeed, we observed no significant changes in sarcomere length, which is in direct contrast to the 2D hiPSC-CM presentation observed by Davis et al.^3^

Functionally, while lactate-purified hiPSC-CMs in 2D monolayer cultures demonstrated aberrant twitch force kinetics and poor calcium handling 5 days post purification (similar to what has been found in cardiomyocytes isolated from ischemic human hearts)^59–61^, our lactate hiPSC-ECTs demonstrated little variability from our MACS hiPSC-ECTs after four weeks in culture. Unlike stress or ischemic heart tissue, the lactate hiPSC-ECTs did not display dysfunction in β-adrenergic responsiveness.^61^ Most importantly, we demonstrate that lactate-generated hiPSC-ECTs retain full contractile function. This ability to directly measure force and contractile kinetics in an integrated human tissue is a great advantage of 3D hiPSC-CM models. By demonstrating an unaltered phenotype between lactate- and MACS-purified hiPSC-ECTs, we ensure there is no bias in our 3D models when testing multiple aspects of function regardless of how the preceding 2D hiPSC-CM purification was performed. It is important to point out the observed increased variability in the functional measures of the ECT groups. While the measured variability does not result in a significant difference between lactate and MACS-purified groups, individual construct heterogeneity may be an experimental design factor to consider when choosing a 2D hiPSC-CM purification method.

An important finding of our study is the global proteome was not significantly impacted by purification method. Changes indicative of an ischemic phenotype, which often include decreased SERCA2a expression, reduction of mitochondrial- and metabolism-related protein pathways, and cardiomyocyte cytoskeletal changes, were not observed in the lactate-purified hiPSC-ECT.^3,59,62,63^ Differentially expressed individual proteins accounted for 0.4% of the proteome identified and did not suggest alterations in any major protein pathways known to be related to ischemia. Indeed, one of the advantages in using 3D hiPSC-ECTs includes molecular signatures of a more mature physiological state.^13^ By demonstrating there is no molecular bias in our lactate hiPSC-ECTs, we ensure this advantage to the 3D model remains. Previous studies in both 2D and 3D hiPSC-CM models have indicated both global protein expression and sarcomere phosphorylation as key markers for maturation^12,13,51^. These data imply lactate purification does not alter the maturation or differentiation state of hiPSC-ECTs.

Additionally, myofilament proteoforms, including PTMs and isoforms, are unaltered between the hiPSC-ECT groups. Both sarcomere phosphorylation and isoform expression for key proteins was found to be similar between lactate and MACS hiPSC-ECTs. It has been shown bottom-up proteomics analyses which digests proteins into peptides prior to MS analysis can obscure post-translational and protein isoform information due to the peptide-to-protein inference problem^41,53^. We have employed top-down proteomics which is a powerful technology for detection and quantification of PTMs and isoforms simultaneously in one spectrum.^13,51,53^ Notably, the atrial form of MLC-2 (MYL7) was significantly increased in the MACS hiPSC-ECTs in our global bottom-up experiments whereas in our top-down data set, MLC-2a is not detected at all. Given the sensitivity and accuracy of our top-down platform and the important role myosin light chain has in development and maturation, this result emphasizes the importance of performing intact protein measurements.^13,55,64^ Here, we have shown there are no alteration in the sarcomeric protein PTMs and isoforms between lactate and MACS hiPSC-ECTs, which implies similar baseline molecular state for the sarcomere.

Taken together, these data indicate there is no difference in function or structure between lactate and MACS hiPSC-ECTs. Whether the absence of the ischemic phenotype observed by Davis et al. in hiPSC-CM in our study is due to the 3D culture configuration, extended time in culture, or both, is beyond the scope of this study. It should be noted the 2D hiPSC-CMs experiments performed by Davis et al. were at most 5 days post-selection and therefore, may have not allowed for sufficient recovery of the hiPSC-CMs after lactate purification. Indeed, while consistency in differentiation, purification method, and inclusion of appropriate controls are all critical within an experimental design, the method of purification does not have a significant impact when hiPSC-CMs are subsequently grown in ECTs.

## Methods

### hiPSC culture

Control hiPSCs were graciously provided by the Stanford Cardiovascular Institute (Stanford University Cardiovascular Institutes Biobank; Hypertrophic Cardiomyopathy/*MYH7*/Control iPSC line)^4^. All hiPSCs were maintained in StemFlex media (Gibco) according to the manufacturer’s protocol. Cryopreserved hiPSCs were thawed and added to StemFlex media supplemented with 5□µM Y-27632 (BD Biosciences). hiPSCs were then plated onto Matrigel (GFR, BD Biosciences) coated six-well dishes. hiPSCs were subsequently cultured at 37°C at 5% CO_2_ until they were 70%–90% confluent with daily media changes before passaging. For passaging, hiPSCs were dissociated using Versene (Gibco), resuspended in StemFlex media, and re-plated onto Matrigel (Corning)-coated plates.

### Differentiation of Human iPSCs into Cardiomyocytes

Human iPSCs from the SCVI wild-type line were differentiated into CMs using a small molecule-directed protocol as previously described^42^. In short, hiPSCs in culture were dissociated and seeded onto Matrigel-coated six-well plates at 1.5–2.0□×□10^6^ cells/well in StemFlex media. Cells were cultured for approximately 3–5□days in StemFlex media [1□day post-100% confluence, at which time differentiation was initiated (day 0)]. On day 0, StemFlex media was replaced with RPMI supplemented with B27 without insulin (Gibco) with a 9□µM CHIR99021 addition (Tocris Bioscience). Exactly 24□hours later (day 1), media was changed to 3□mL/well RPMI + B27 without insulin and cells were cultured in this media for 48□h (day 3). On day 3, the media was changed to 3□mL/well RPMI + B27 without insulin supplemented with 5□μM IWP-2 (Stemgent). Forty-eight hours later (day 5), the media was changed to 3□mL RPMI + B27 without insulin. The media was changed to RPMI + B27 complete supplement (with insulin) (Gibco) on day 7, and the differentiated cells were maintained in this media until day 15 with media changes every 48–72□h. On day 15, cells from wells containing ≥80% contracting cells by visual inspection were dissociated with 10× TrypLE (Thermo Fisher Scientific) according to the manufacturer’s protocol. Cells were then cryopreserved using a 1:9 ratio of DMSO (Millipore Sigma) and FBS (Thermo Fisher Scientific).

### hiPSC-CM purification

After thawing and suspending in EB20 media^2^, cells were replated on Synthemax (Corning) coated six-well plates. Each hiPSC-CM batch (5 batches of 15 million cells) was matched for both lactate and MACS purifications. For the lactate purified hiPSC-CMs, 48 hours after replating, hiPSC-CMs were purified using CDM3L media, made with RPMI 1640 no glucose (Life Technologies), 500 μg/mL recombinant human albumin, 213 μg/mL l-ascorbic acid 2-phosphate, and 4□mM l-lactic acid (Sigma-Aldrich)^16^ for 7 days with media changes every 48–72□h. Following selection, hiPSC-CMs were maintained in RPMI with B27 supplement until day 30 at which point hiPSC-CMs were combined with hiPSC-CFs to generate ECTs. For the MACS selected cells, thawed cells were maintained in RPMI with B27 supplement until day 30, or the day of ECT generation. Cells were dissociated from the plate using 10x TrypLE and purified with the MACS system (Miltenyi Biotec) according to the manufacturer’s protocol. In sum, the cell suspension was quenched with EB20, passed through a 100 µM cell strainer (Fisher Scientific), counted, and centrifuged. The remaining cell pellet was then resuspended 80µL/5x10^6^ cells in “Miltenyi” buffer containing 0.5% w/v BSA (Millipore Sigma), DPBS, and 2 mM EDTA (Fisher Scientific). Non-Cardiomyocyte Depletion Cocktail (Miltenyi Biotec) was added in a 20µL/5x10^6^ cells and incubated at 4°C for 5 minutes. The mixture was then washed with additional “Miltenyi” buffer and centrifuged. The cell pellet was then resuspended 80µL/5x10^6^ cells in “Miltenyi” buffer, and Anti-Biotin MicroBeads (Miltenyi Biotec) were added in a 20µL/5x10^6^ cells ratio and incubated at 4°C for 10 minutes. Additional buffer was added to bring final concentration to 500 µL/5x10^6^ cells. 500 µL of the cell suspension was loaded into the LS Column/MACS Separation apparatus (Miltenyi Biotec). Flow through of the pure hiPSC-CMs was centrifuged, resuspended, and taken for either ECT generation or flow cytometry analysis.

### Differentiation of Human iPSCs into Fibroblasts

Isogenic hiPSC-cardiac fibroblasts (CFs) were generated as previously described.^43^ In brief, hiPSCs were dissociated with Versene seeded on Matrigel coated 6-well plates at the density of 1.5–2.0□×□10^6^ cells/well in mTeSR1 media (WiCell) supplemented with 10□μM Y-27632. Cells were maintained in mTeSR1 media for approximately 5 days with daily changes until they reached 100% confluency (day 0). On day 0, the cells were treated with 2□ml RPMI+B27 without insulin and supplemented with 12□µM CHIR99021 for 24□h. After 24 hours, the media was changed to 2□ml RPMI+B27 without insulin for 24□h. After 24 hours, the media was changed to 2□ml of the CFBM media with 75□ng/ml bFGF (WiCell). Subsequently, cells were maintained with CFBM+75□ng/ml bFGF every other day until day 20. On day 20, cells were either taken for flow cytometry analysis or cryopreserved using a 1:9 ratio of DMSO (Millipore Sigma) and FBS (Thermo Fisher Scientific). Once thawed, the hiPSC-CFs were maintained in FibroGRO-LS media (Millipore Sigma) in uncoated six-well culture plates (Corning) and passaged every 5-6 days. Low passage number (<12) were used for hiPSC-ECT generation.

### Flow Cytometry

hiPSC-CMs and hiPSC-CFs were analyzed as previously described^42,43^. Briefly, dissociated cells were vortexed to disrupt the aggregates followed by neutralization by adding equal volume of EB20 media. Cells were counted to designate 1 million cells for labeling. Cells were fixed in 1% paraformaldehyde, washed with FACS buffer (DPBS, 0.5% bovine serum albumin (BSA), 0.1% NaN3), centrifuged and resuspended in about 50□μl FACS. Primary antibodies, including monoclonal anti-α-actinin (IgG1, Sigma, 1:500 dilution) and monoclonal anti-cTnT (IgG1, Thermo Scientific, 1:200 dilution), were incubated with the cells according to the manufacturer’s instructions. Afterwards, cells were washed with FACS buffer plus 0.1% Triton X-100, centrifuged, and all but 50□μl supernatant discarded. Secondary antibody appropriate to the primary IgG isotype was diluted at 1:1000 in FACS buffer plus 0.1% Triton X-100. Samples were incubated for 30 minutes at room temperature, washed in FACS buffer, and resuspended in FACS buffer for analysis. Data were collected on a FACSCalibur (Beckton Dickinson) and Attune Nxt (ThermoFisher) flow cytometers and analyzed using FlowJo.

### hiPSC-ECT Generation

Day 30 lactate-purified hiPSC-CMs were dissociated with 10x TrypLE and counted using a hemocytometer. MACS-purified hiPSC-CMs were taken straight from purification and counted using hemocytometry. hiPSC-CMs were subsequently resuspended at 2×10^6^ CM/mL in fibrin ECT media (60.3% high-glucose DMEM; 20% F12 nutrient supplement; 1□mg/mL gentamicin; 8.75% FBS; 6.25% horse serum; 1% HEPES; 1× nonessential amino acid cocktail; 3□mM sodium pyruvate; 0.004% (wt/vol) NaHCO3; 1□µg/mL insulin; 400□µM tranexamic acid; and 17.5□µg/mL aprotinin)^13^ and incubated for at least 1.5h on a rotating platform at 37°C. Low-passage isogenic hiPSC-CFs were dissociated using 1×TrypLE (Thermo Fisher Scientific) and counted using a hemocytometer. After rotational culture, hiPSC-CMs were mixed with hiPSC-CFs in a 10:1 ratio per hiPSC-ECT, as previously described.^65^ 1.25□mg/mL fibrinogen and 0.5 unit of thrombin were added to the cell mixture, and the cell suspension was quickly loaded onto a 20 × 3-mm cylindrical mold of FlexCell Tissue Train silicone membrane culture plate. Following polymerization of the fibrin matrix, ECTs were maintained with fibrin ECT media at 37°C with 5% CO_2_ for 4 weeks with media changes every 2– 3□days.

### Twitch Force and Ca^2+^-Transient Measurements

Isometric twitch force and Ca^2+^-transients were measured in hiPSC-ECT using protocols similar to those previously described.^13,66^ In brief, each hiPSC-ECT construct was attached using sutures to a model 801B small intact fiber test apparatus (Aurora Scientific) in Krebs–Henseleit buffer [119□mmol/L NaCl, 12□mmol/L glucose, 4.6□mmol/L KCl, 25□mmol/L NaHCO3, 1.2□mmol/L KH2PO4, 1.2□mmol/L MgCl2, 1.8□mmol/L CaCl2, gassed with 95% O2-5% CO2 (pH 7.4)] at 37°C. Krebs–Henseleit buffer flowed throughout the experiments at a rate of 1□mL/min. Force readouts were performed on a model 403A force transducer (Aurora Scientific). Stimulation was initiated on each hiPSC-ECT at 1□Hz (2.5□ms, 12.5□V). Each construct was stretched to optimal length (until maximal twitch force was achieved). Constructs were left to equilibrate for 20□min at 1 Hz. Following equilibration, twitch force production was measured with pacing at a frequency of 1.5□Hz both at baseline and following 5□min preincubation with 1□µM isoproterenol. Automaticity was captured after pacing. After functional analyses, hiPSC-ECTs were washed with DPBS, flash frozen, and stored at - 80°C until proteomic analysis.

hiPSC-ECTs were then introduced to a Fura-2 loading solution consisting of Krebs–Henseleit buffer supplemented with 5□μM Fura2-AM (Invitrogen) and 1% (vol/vol) Chremophor EL (Sigma) with constant oxygenation (95% O2, 5% CO2) for 30□min at 37°C. Following Fura-2 loading, ECTs were left to equilibrate for 40□min with perfusion with Krebs–Henseleit buffer at a rate of 1□mL/min and paced at 1□Hz. Both twitch force and calcium transient data were recorded at pacing frequency 1.5 Hz. Fura-2 fluorescence was measured by alternately illuminating the preparation with 340-and 380-nm light (at a frequency of 250□Hz) while measuring the emission at 510□nm using IonOptix hardware and software (IonOptix Corporation, Milton, MA). The emitted fluorescence and force data were stored as the 340- and 380-nm counts and as the ratio R = F340/F380. Data were analyzed using IonWizard 6.0 software (IonOptix). Under each condition, 40–60 successive contractions were collected and averaged. These data were exported to Microsoft Excel for parameter calculations. Statistical significances were determined by normal t-test with α = 0.05 with two-sided analysis. All data represented as data mean ± SEM.

### Cryopreservation, Sectioning, and Immunohistochemistry

hiPSC-ECTs were rinsed in DPBS (Thermo Fisher Scientific) in well and incubated in 30□mM 2,3-butanedione monoxime in DPBS for 5□min. hiPSC-ECTs were then exposed to a filtered 30% sucrose (in DPBS) solution for 1 h at room temperature, followed by a 1-h incubation in a 1:1 mixture of optimal cutting temperature (OCT) compound (Tissue-Tek) and 30% sucrose solution in DPBS. The sucrose solution was aspirated from the well and hiPSC-ECTs were covered with OCT compound in well before freezing the wells on a metal plate on dry ice. Cryopreserved hiPSC-ECTs were then stored at −80°C. The cryopreserved hiPSC-ECT disks were taken out of the plate before sectioning. Cryopreserved hiPSC-ECTs were then sectioned lengthwise at 6-µm thickness (Leica CM 1950UV), mounted onto charged slides (Superfrost +, Fisherbrand), and fixed in 100% acetone for 15□min at 4°C. After drying, slides were placed in a vertical washer and rinsed with water for 10□min. Slides were then rehydrated in PBS and incubated in blocking buffer [0.15% Triton-X-100, 5% normal goat serum (NGS), 2 mg/mL BSA in PBS] for 1□h at room temperature. Sections were incubated with primary antibodies overnight in a humidified chamber at 4°C with α-actinin (1:1,000, Sigma, A7811). Slides were incubated with secondary antibody (Alexa Fluor Plus 488; Invitrogen) at 5–8 µg/mL in blocking buffer for 1□h at room temperature in a humidified chamber. Following labeling, sections were cover slipped using Prolong Gold Antifade Reagent (Invitrogen) with 4′,6-diamidino-2-phenylindole (DAPI) to label nuclei. Imaging was performed using Leica SP8 Confocal WLL STED Microscope using the 100x objective and Leica imaging software. Multiple sections from each hiPSC-ECT were imaged and 100 sarcomeres were measured using manual annotation on the Leica imaging software. Statistical significances were determined by normal t-test with α = 0.05 with two-sided analysis. All data represented as data mean ± SEM.

### Global Bottom-up Proteomics

Global proteomics was performed similarly as previously described.^67^ Samples were randomized and blindly prepared to avoid bias. Cryopreserved hiPSC-ECTs were gently thawed on ice and homogenized in 40 μL of Azo^46^ buffer (0.1% w/v 4-hexylphenylazosulfonate (Azo), 25 mM ammonium bicarbonate, 10 mM l-methionine, 1 mM dithiothreitol (DTT), and 1× HALT protease and phosphatase inhibitor) using a handheld Teflon homogenizer (Thomas Scientific). Samples were centrifuged, and the supernatant was normalized to 1 mg/mL using 0.1% Azo buffer by the Bradford assay (Bio-Rad). Samples were reduced with 30 mM DTT for 1 h and subsequently alkylated with 30 mM iodoacetamide for 1 h. Trypsin Gold (Promega) was added in a 50:1 ratio to the samples and incubated for 24 h. After 24 h, trypsin was quenched with formic acid. Azo was then degraded at 305 nm for 5 min (Analytik Jena). Samples were centrifuged, and the resulting supernatant was desalted using 100 μL Pierce C18 tips (Thermo Fisher Scientific) according to the manufacturer’s protocol. The peptides were dried in a vacuum centrifuge and resuspended in 0.1% FA. The peptide concentrations were determined using Nanodrop One Microvolume UV-Vis Spectrophotometer (ThermoFisher).

LC-TIMS-MS/MS was performed using a nanoElute nanoflow LC system (Bruker Daltonics) coupled to the timsTOF Pro (Bruker Daltonics). 200 ng of each peptide sample was loaded on an Aurora Elite capillary C18 column (IonOpticks). Peptides were separated using a 120 min gradient at a flow rate of 400 nL/min (mobile phase A (MPA): 0.1% FA; mobile phase B (MPB): 0.1% FA in acetonitrile). For the first 60 minutes, a gradient of 2–17% MPB was applied, then 17 to 25% MPB for the next 30 min, 25–37% MPB for 10 minutes, 37–85% MPB for 10 minutes, and ending at 85% MPB for an additional 10 min. The column utilized nanoESI source for sample passage to the mass spectrometer. MS spectra were captured with a Bruker timsTOF Pro quadrupole-time of flight (Q-TOF) mass spectrometer (Bruker Daltonics, Billerica, MA, USA) operating in diaPASEF mode, using 32 windows ranging from m/z 400 to 1200 and 1/K_0_ 0.6 to 1.42. Data processing occurred similar as previously described.^68^ LC-MS data were processed using DIA-Neural Network (DIA-NN)^69^ using the default parameters unless noted in the following: 1% FDR, Library-free search enabled, Minimum fragment m/z: 200, Maximum fragment m/z: 1800, Minimum precursor m/z: 400, Maximum precursor m/z: 1200, Minimum precursor charge: 2, Maximum precursor charge: 4, Minimum peptide length: 7, maximum peptide length: 30, Maximum missed cleavages: 2, MS1/MS2 mass accuracy: 10 ppm, Quantification strategy: Robust LC (High Precision), Neural network classifier: Double-pass mode. All other analysis was performed in R. Protein-level quantification data were filtered using the “DAPAR” package^70^ to include proteins identified in 2 of 3 runs in at least one sample group. Values were then median normalized and missing values were imputed via ssla for partially observed values within a condition or set to the 2.5% quantile of observed intensities for observations that were missing entirely within a condition. The “DEP” R package was used to perform a Limma test between all specified contrasts, and the “IHW” R package was used to adjust all p-values, using the number of quantified peptides per protein as a covariate. A p_adj_ threshold of 0.05 and a log_2_ Fold Change threshold of 0.6 were set to identify significant changes to protein abundance. Subsequent data were visualized and plotted using “ggplot2”^71^. All data represented as data mean ± SEM.

### Myofilament Top-down Proteomics

Myofilament proteomics was performed similarly as previously described.^51^ All steps were kept at 4 °C to minimize artifactual modifications.^52^ In sum, cryopreserved hiPSC-ECTs were gently thawed on ice and homogenized in 50 μL of HEPES extraction buffer (25 mM HEPES (pH 7.4), 60 mM NaF, 1 mM l-methionine, 1 mM DTT, 1 mM PMSF in isopropanol, 1 mM Na3VO4 containing protease and phosphatase inhibitors) using a handheld Teflon homogenizer. The cell homogenate was centrifuged, and the supernatant containing cytosolic proteins was discarded. The resulting pellet was washed again in HEPES extraction buffer, centrifuged, and supernatant discarded. The pellet was then homogenized in 40 μL of TFA extraction buffer (1% TFA, 5 mM TCEP, 5 mM l-methionine) using a handheld Teflon homogenizer. The homogenate was centrifuged, and the resulting supernatant was desalted with five volumes of MPA (0.1% formic acid in HPLC grade water) using a 10 kDa molecular weight cutoff filter (Amicon). Protein normalization was performed using a bovine serum standard curve and Bradford assay.

A NanoAcquity LC system (Waters) was used as part of the reverse phase chromatography (RPC) system. 500 ng of the protein extracts were run through a home-packed PLRP-S capillary column (200 mm long, 0.25 mm i.d., 5 μm particle size, 1000 Å pore size; Agilent Technology). The column was heated to 60 °C at an 8 μL/min flow rate. The gradient is as follows, in terms of MPB (0.1% formic acid in 50:50 acetonitrile and ethanol): 10% MPB at 0-5 min, slow increase to 65% at 5-65 min, 95% at 70-75 min, back to 10% at 75.1 min, and steadily at 10% until 80 min. The eluted proteins were analyzed using a Bruker Impact II quadrupole-time-of-flight (Q-TOF) mass spectrometer (Bruker). The mass spectrometer parameters are as follows: end plate offset – 500 V; capillary voltage – 4500 V; nebulizer – 0.3 bar; dry gas flow rate – 4.0 L/min at 200 °C; quadrupole low mass – 650 m/z; scan rate – 1 Hz; m/z range – 200– 3000 m/z. Three technical replicates were collected for one sample across instrument run time to ensure data reproducibility and stability.

DataAnalysis 4.3 software (Bruker Daltonics) was used for all MS data analysis. A smoothing width of 2.01 using the Gaussian algorithm was applied to each chromatogram. The Maximum Entropy algorithm within the DataAnalysis 4.3 software was implemented to deconvolute spectra for proteins of interest at resolving power of 50,000. The sophisticated numerical annotation procedure (SNAP) algorithm was used to determine the monoisotopic masses of all deconvoluted ions. Relative quantification of protein phosphorylation is reported from deconvoluted spectra, in a relative abundance of a particular proteoform (Ptotal). Ptotal is equivalent as the ratio of the peak intensity of the proteoform (mol Pi) to the total sum of peak intensities of all proteoforms (mol protein) of the same protein. Statistical significances were determined by normal t-test with α = 0.05 with two-sided analysis. All data represented as data mean ± SEM.

### Statistical Analyses

All statistical analyses were performed as 2-sided t-tests with assumed normal distribution, unless otherwise stated, with α = 0.05. All data represented as data mean ± SEM, unless otherwise stated.

### Study Approval

Protocols for the generation of hiPSCs were approved by the Stanford University Human Subjects Research Institutional Review Board and written consent was obtained from all study participants.

## Data Availability

Source data for this manuscript are available via the MassIVE repository with identifier ID=a452c86f653c4e4d9144c497b53dc062 (ftp://MSV000091869@massive.ucsd.edu). Functional data is uploaded via Excel Spreadsheet in the supplemental materials. During review, the data are private but still available to reviewers at massive.ucsd.edu with the password “kjr”.

## Author Contributions

KJR, TJK, JCR, and YG contributed to study design. KJR cultured cells upon differentiation, generated hiPSC-ECTs, performed proteomics sample preparation, analyzed targeted proteomics analysis, prepared hiPSC-ECTs for sections, imaged IHC images and processed data, wrote the manuscript. WdL gathered and analyzed all functional data on hiPSC-ECTs. MWM and TJA analyzed global proteomics data. JAM assisted with collection of targeted proteomics data. JZ and GK differentiated cells into CMs and CFs. YZ assisted with global proteomics data collection. TJK, JCR, and YG edited the manuscript and provided supervision of the study. All authors contributed to the writing of the manuscript.

## Acknowledgements

This research is supported by NIH R01 HL096971. YG would like to acknowledge NIH R01 GM125085, R01 HL109810, and S10 OD018475. KJR acknowledges the National Science Foundation Graduate Research Fellowship Program under Grant No. DGE-1747503 and the Graduate School and the Office of the Vice Chancellor for Research and Graduate Education at the University of Wisconsin-Madison, funded by Wisconsin Alumni Research Foundation. K.J.R would like to acknowledge David S. Roberts for his helpful discussion. All authors would like to acknowledge Joseph Wu and the Stanford Cardiovascular Institute for their gracious gift of the hiPSC lines. Some figures made in Biorender.

