## Supplemental Figures for "Lactate and Immunomagnetic-purified iPSC-derived Cardiomyocytes Generate Comparable Engineered Cardiac Tissue Constructs"

**SUPPLEMENTAL TABLE OF CONTENTS**

*Supplemental Figures………………………………………………………………….Page #*

Supplemental Figure 1. Purification Efficiency of Lactate and MACS methods………….3

Supplemental Figure 2. Immunohistochemistry Images for Sarcomere Length Analysis…………………………………………………………………………………………..4

Supplemental Figure 8. Differentially Expressed Proteins between Lactate and MACS hiPSC-ECTs……………………………………………………………………………..……..10

Supplemental Figure 9. Pathway Analysis of hiPSC-ECT proteome……………..……..11

Supplemental Figure 10. Base Peak Chromatograms (BPCs) of Intact Sarcomere Proteomics……………………………………………………………………………………...12

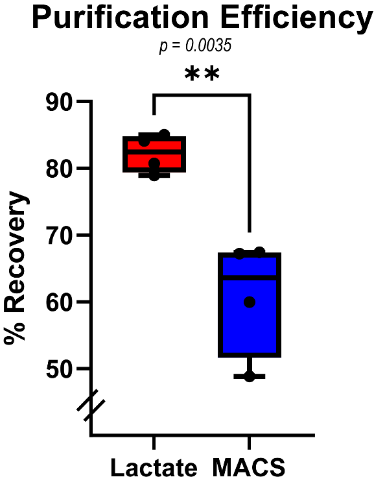

**Supplemental Figure 1. Purification Efficiency of Lactate and MACS methods.** Efficiency of purification methods are given by percentage of pure hiPSC-CMs recovered from matched successful differentiation batches (evaluated visually with greater than 80% of well contracting). Lactate differentiation was significantly more successful in hiPSC-CM recovery in comparison to MACS (lactate = 82.21 ± 2.858 %, vs. MACS = 60.87 ± 8.727 %; p = 0.0035). All tests performed with four separate differentiation batches. Statistical analysis is t test with α = 0.05.

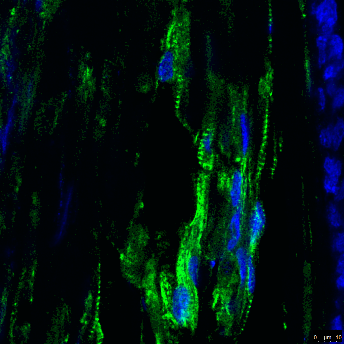

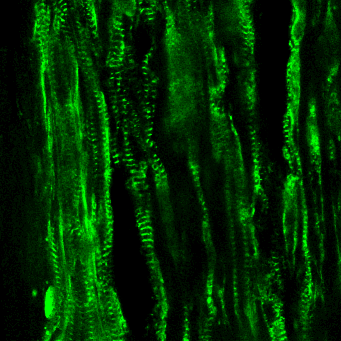

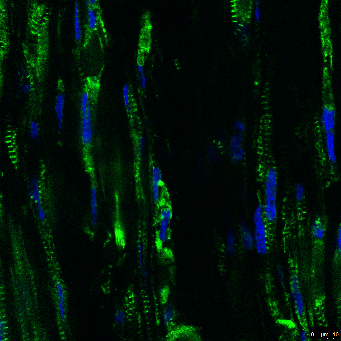

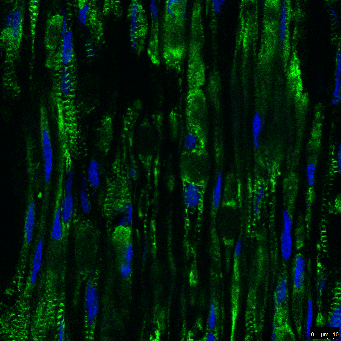

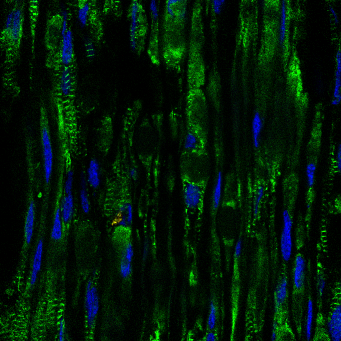

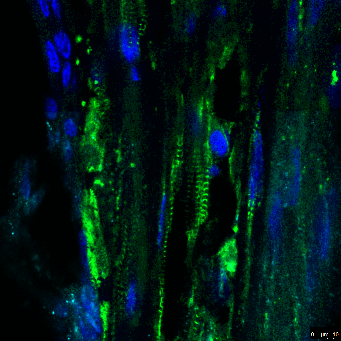

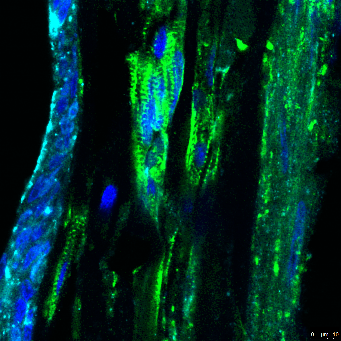

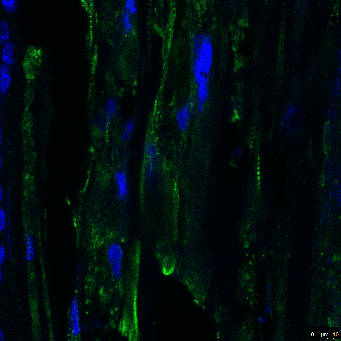

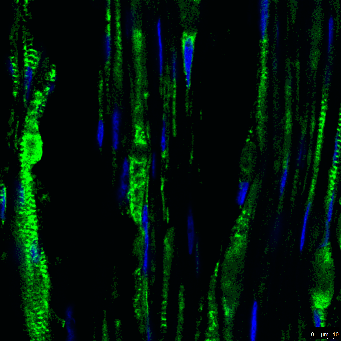

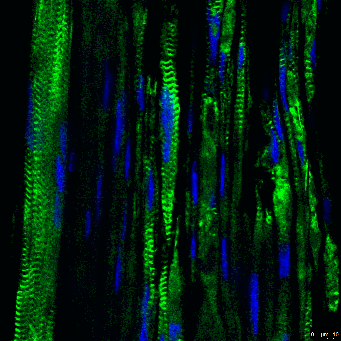

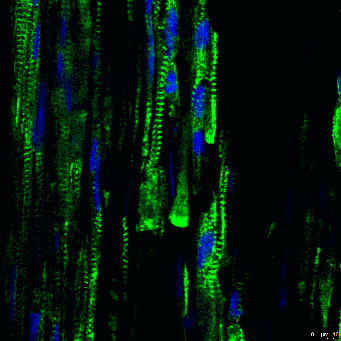

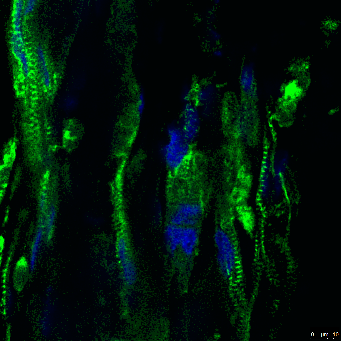

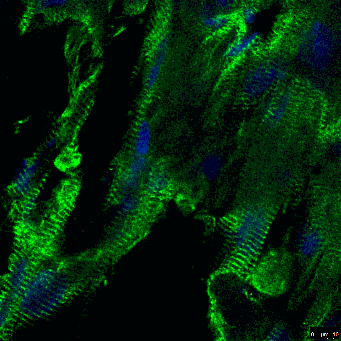

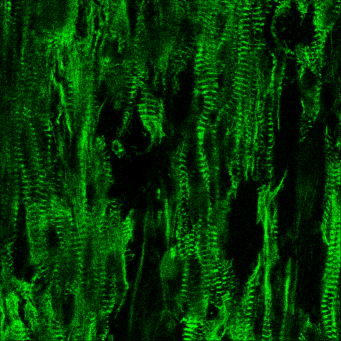

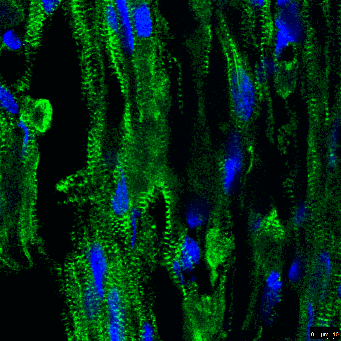

**Supplemental Figure 2. Immunohistochemistry Images for Sarcomere Length Analysis.** *Top (left to right) to Bottom – Condition Replicate Number (Slide Number):* Lactate 1(1), Lactate 2(2), Lactate 2(4), Lactate 2(6), Lactate 2(7), Lactate 3(11), Lactate 3(16), Lactate 3(18), MACS 1(1), MACS 1(4), MACS 1(5), MACS 2(1), MACS 2(2), MACS 3(1), MACS 3(2). Green fluorescence indicates alpha-actinin staining. DAPI is indicated in blue.

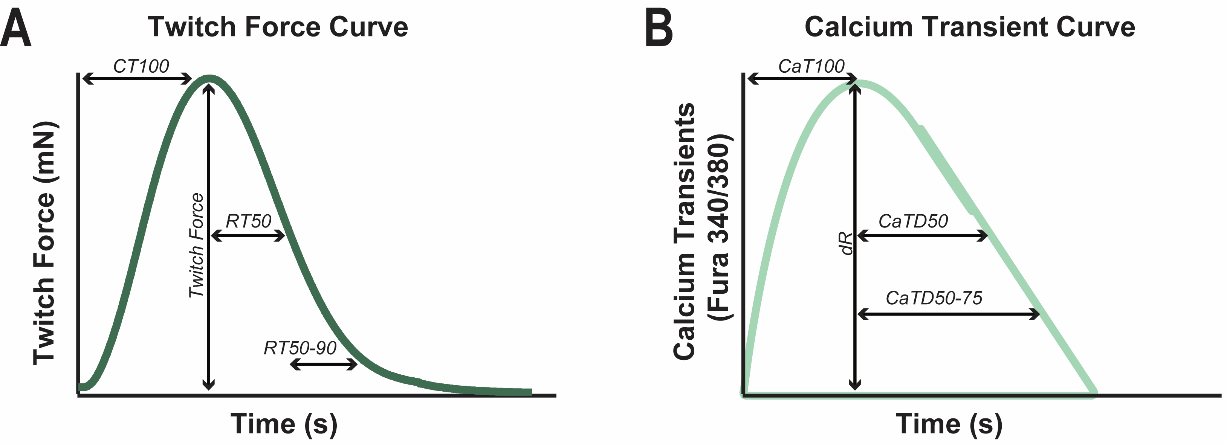

**Supplemental Figure 3. Overview of Functional Parameters.** (A) Twitch force parameters as indicated: twitch force amplitude (TF), time from pacing stimulus to twitch force peak (CT100), time from twitch force peak to 50% twitch force decay (RT50), and time from 50% to 90% twitch force decay (RT50–90). (B) Ca^2+^ transient parameters as indicated: Ca^2+^TR peak (dR), time to Ca^2+^TR peak (CaT_100_), time from Ca^2+^TR peak to 50% Ca^2+^TR decay (CaDT_50_), and time from 50% Ca^2+^TR decay to 75% Ca^2+^TR decay (CaDT_50-75_).

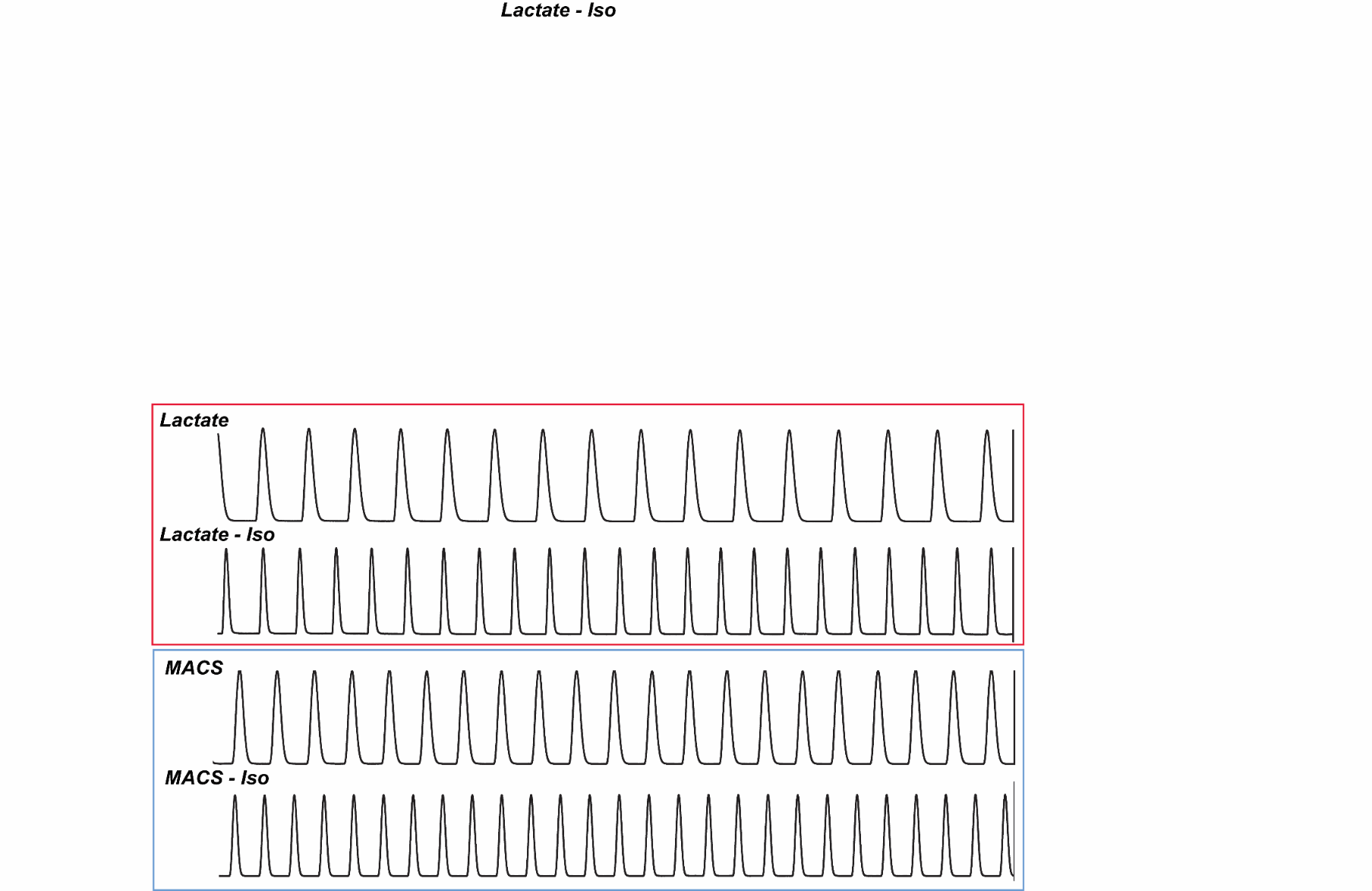

**Supplemental Figure 4. Representative Automaticity Traces of MACS and Lactate hiPSC-ECTs.** Traces representative of automaticity in lactate (red) and MACS (blue) hiPSC-ECTs for approximately 15 seconds after functional evaluation. Traces indicate no arrythmias in automaticity pre- or post-isoproterenol treatment.

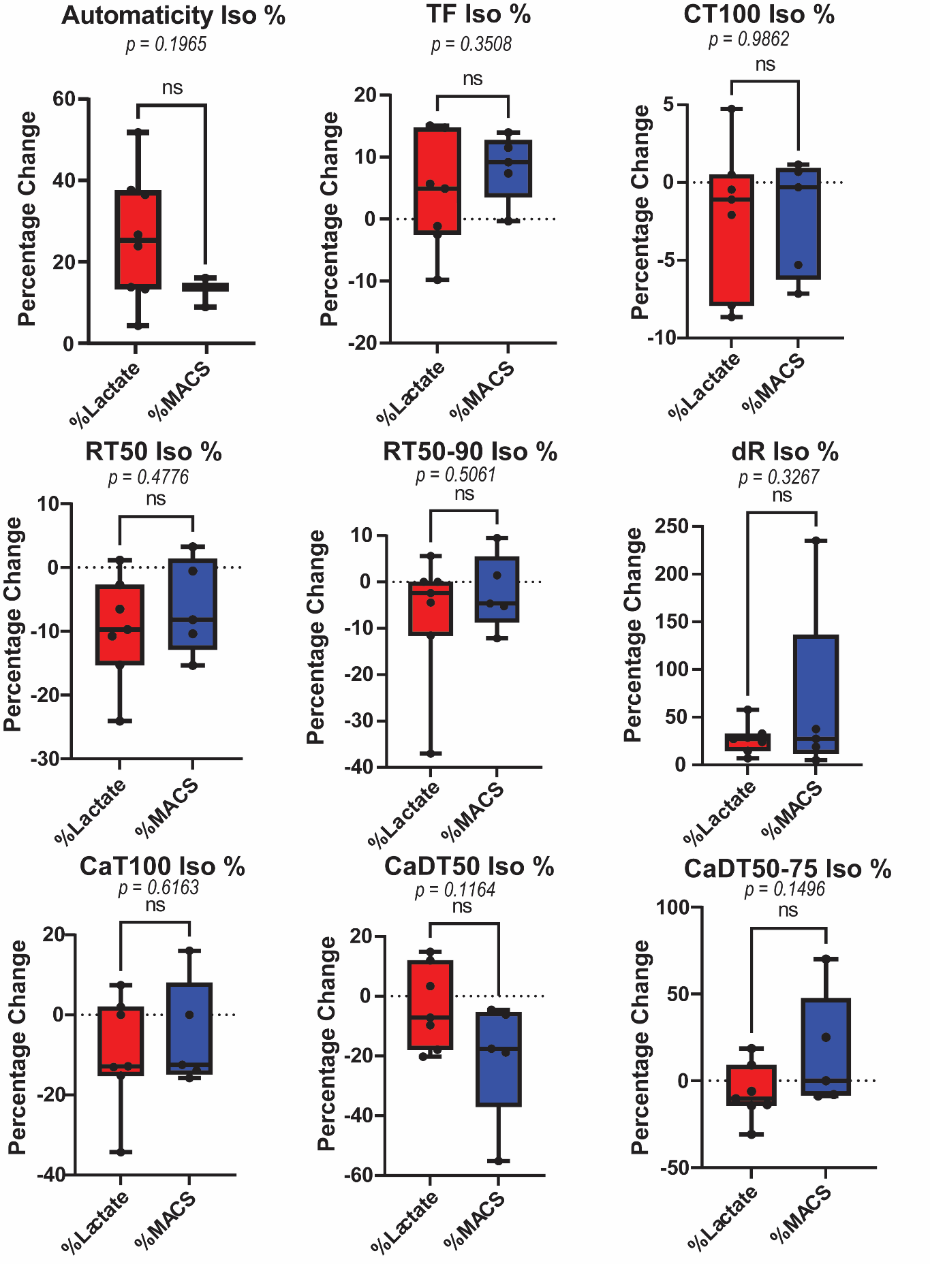

**Supplemental Figure 5. Isoproterenol Assessment of hiPSC-ECTs.** Functional parameters given as a percentage change in response to isoproterenol: automaticity (lactate = 27.28 ± 5.870 %, n=8 vs. MACS = 13.35 ± 2.237 %, n=5, p = 0.1965); TF (lactate = 3.847 ± 3.4440 %, vs. MACS = 8.345 ± 2.4270 %, p = 0.3508); CT_100_ (lactate = -2.136 ± 1.7850 %, vs. MACS = -2.181 ± 1.6900 %, p = 0.9862); RT_50_ (lactate = -9.700 ± 3.1440 %, vs. MACS = -6.239 ± 3.3670 %, p = 0.4776); RT_50-90_ (lactate = -7.117 ± 5.2590 %, vs. MACS = -2.222 ± 3.6270, p = 0.5061); dR (lactate = 27.430 ± 6.0710 % vs. MACS = 64.840 ± 42.9200 %, p = 0.3267); CaT_100_ (lactate = -9.405 ± 5.2970 %, vs. MACS = -5.236 ± 5.9890 %, p = 0.6163); CaDT_50_ (lactate = -3.540 ± 5.2560 %, vs. MACS = -20.470 ± 9.1410 %, p = 0.1164); CaDT_50-75_ (lactate = -6.776 ± 6.1460 %, vs. MACS = 15.660 ± 14.9000 %, p = 0.1496). All tests performed with biological replicates as lactate, n=7 and MACS, n=5, unless otherwise stated. All statistical analyses are unpaired t tests with α = 0.05.

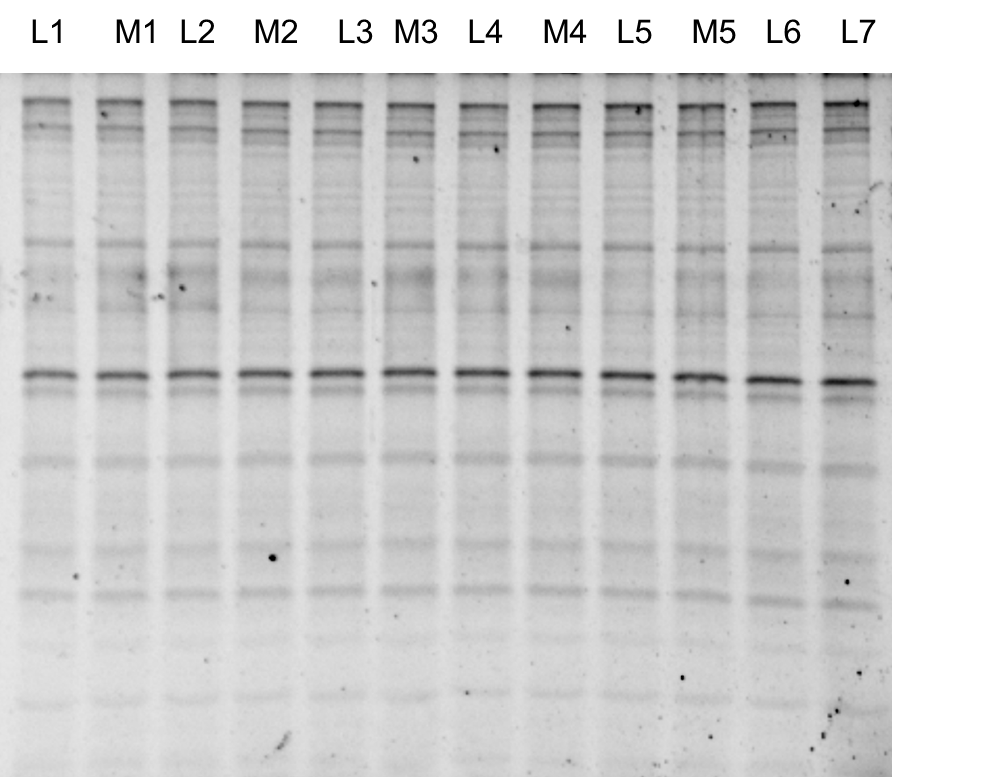

**Supplemental Figure 6. SDS-Page Gel for Reproducibility of Proteomics Extraction.** 12.5% SDS-Page gel with lactate (L#) and MACS (M#) replicates. 500 ng of protein lysate was loaded onto the gel.

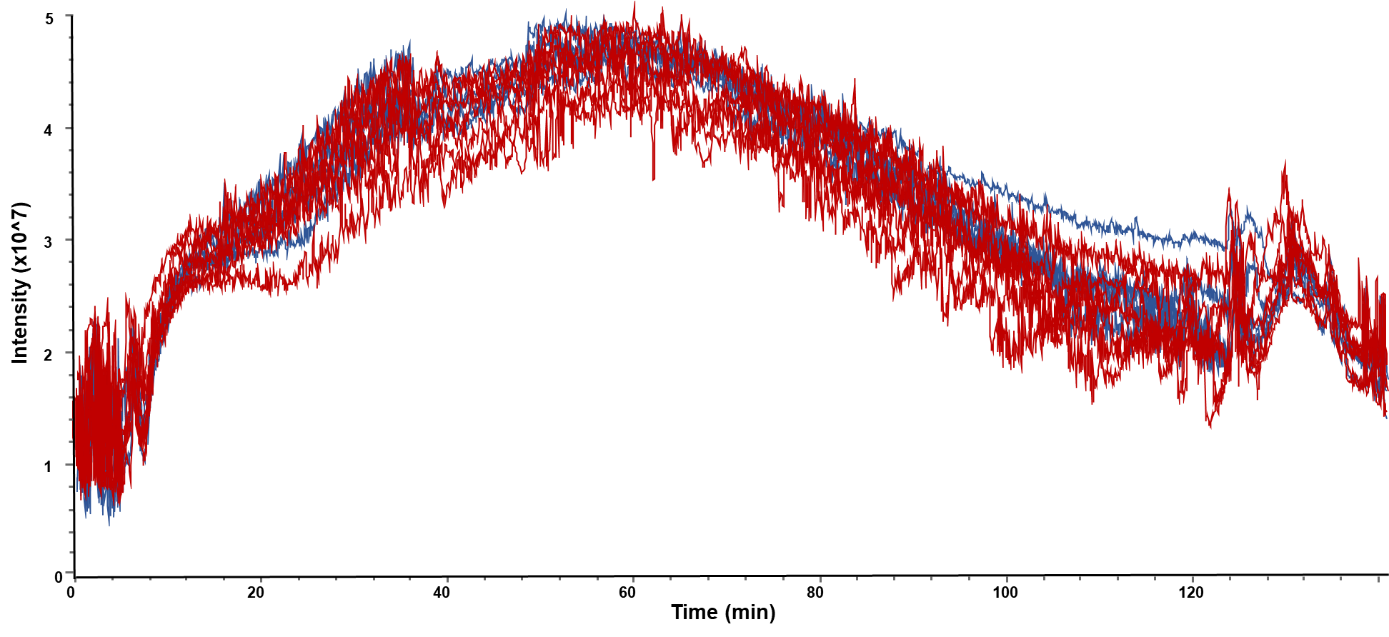

**Supplemental Figure 7. Total Ion Chromatograms (TICs) for Global Proteomics.** TIC traces are shown for lactate (red) and MACS (blue) biological replicates. Overlay of the TICs indicates good reproducibility for all samples.

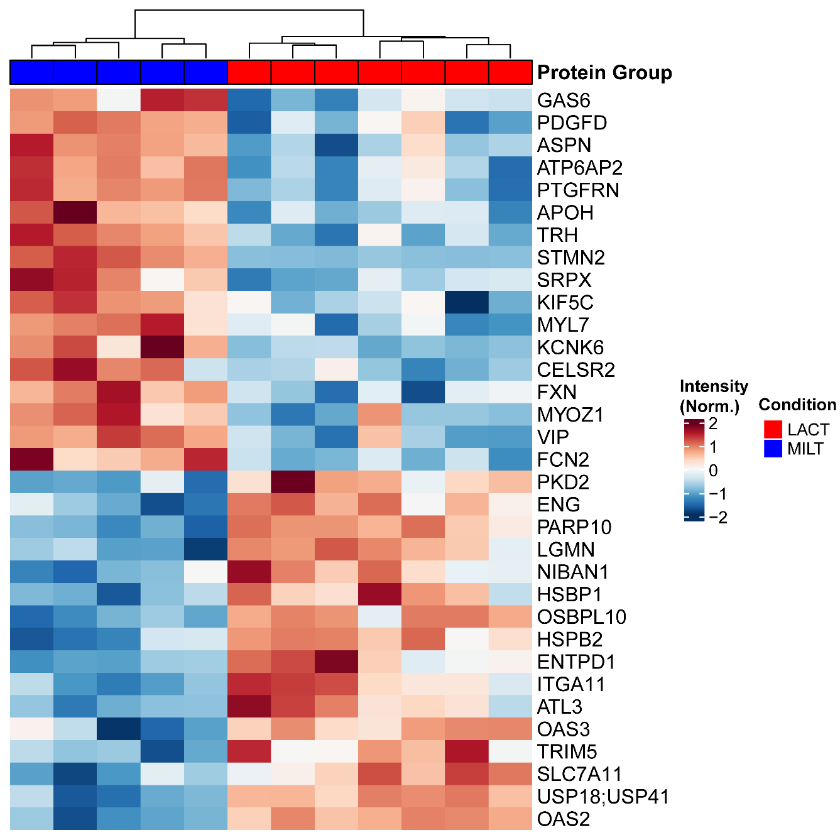

**Supplemental Figure 8. Differentially Expressed Proteins between Lactate and MACS hiPSC-ECTs.** Heatmap depicts normalized intensity values between differentially expressed proteins.

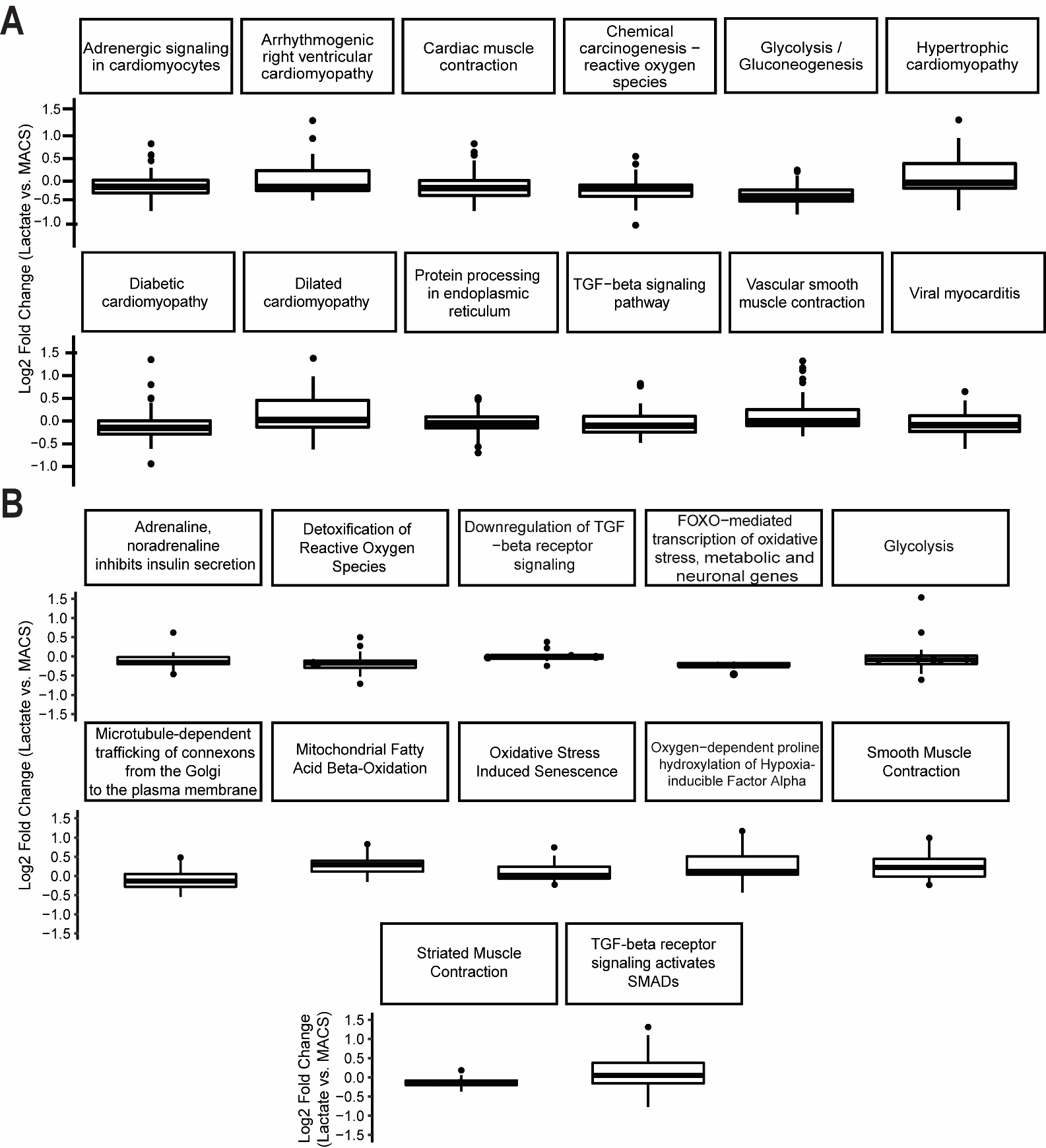

**Supplemental Figure 9. Pathway Analysis of hiPSC-ECT proteome.** (A) KEGG pathway analysis for global protein expression. (B) UniProt pathway analysis for global protein expression.

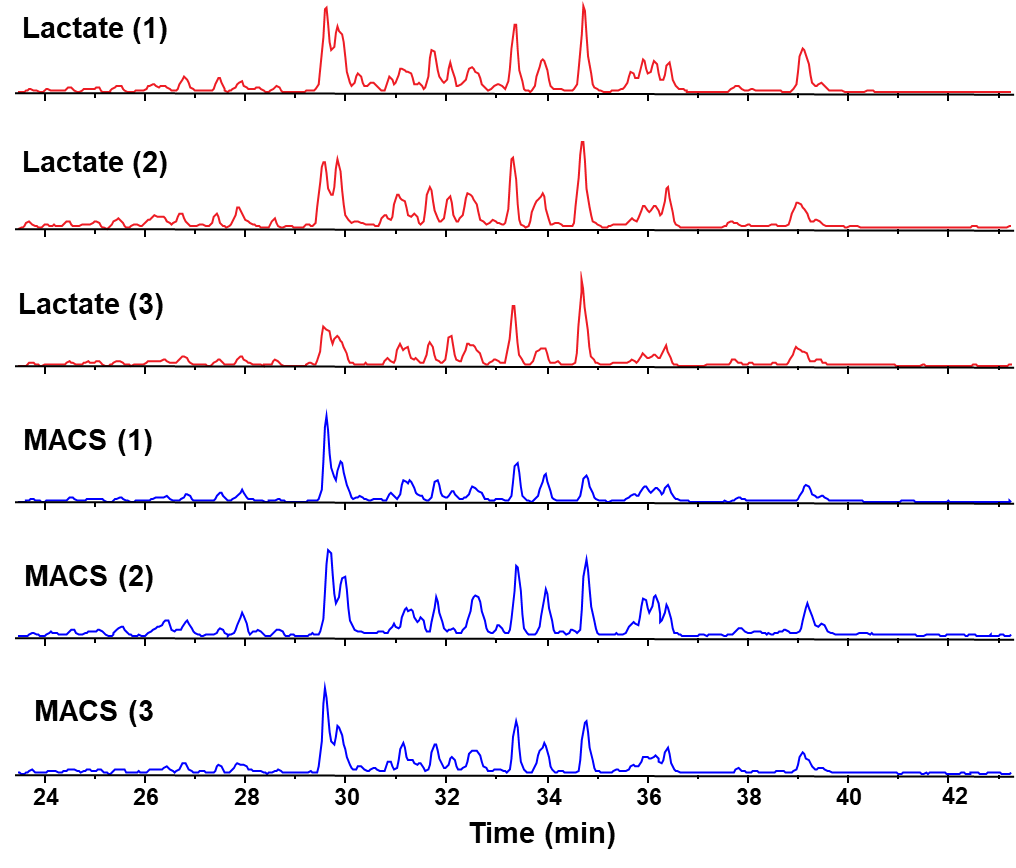

**Supplemental Figure 10. Base Peak Chromatograms (BPCs) of Intact Sarcomere Proteomics.** BPC traces are shown for representative lactate (red) and MACS (blue) biological replicates with normalized intensity. Similar intensity and BPC shape indicates good reproducibility for all samples.

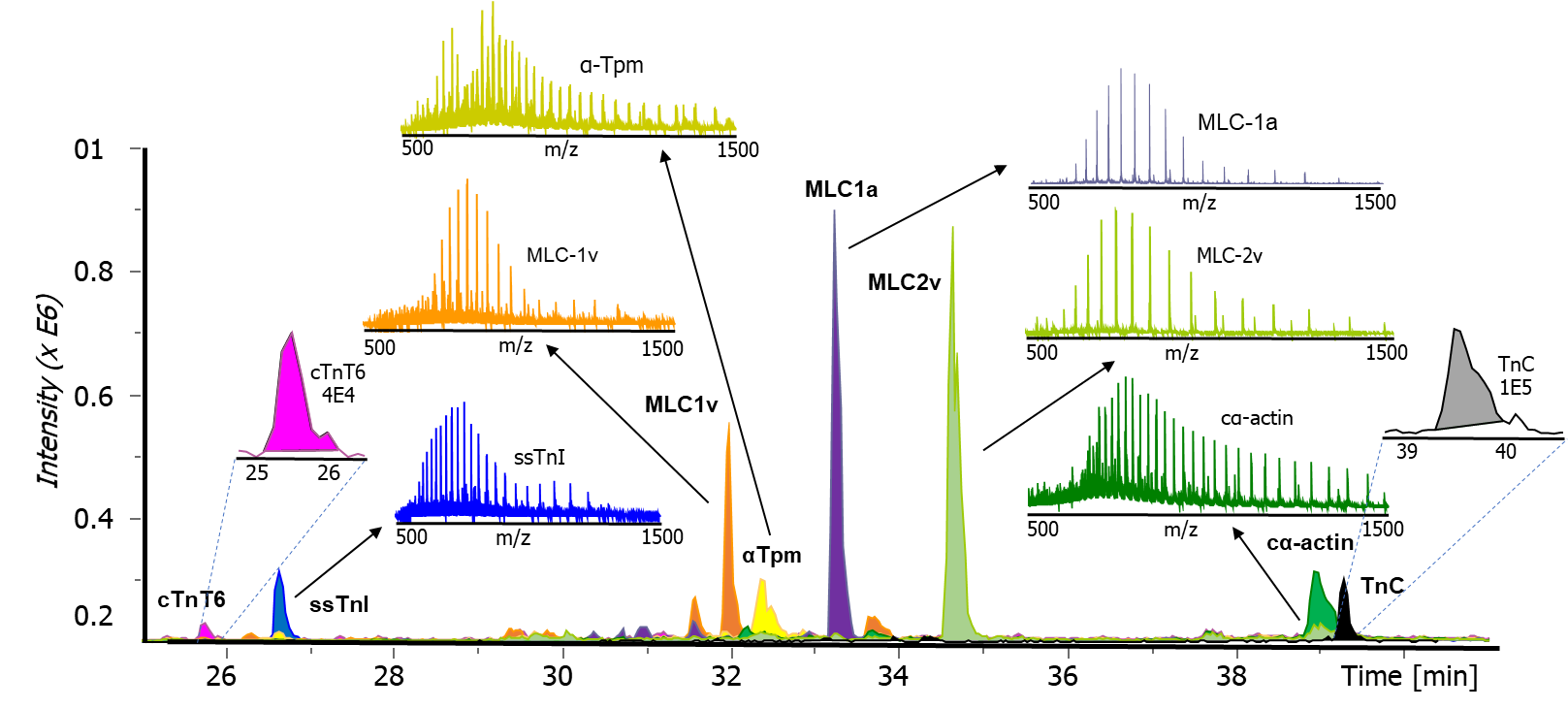

**Supplemental Figure 11. Myofilament Proteins in Intact Protein Mass Spectrometry Analysis.** Contractile proteins such as cardiac troponin T (cTnT6), slow-skeletal troponin I (ssTnI), ventricular myosin light chains (MLC-1v/MLC-2v), alpha tropomyosin (α-Tpm), atrial myosin light chain (MLC-1v), cardiac alpha-actin (cα-actin), and troponin C (TnC) are all identifiable from a 500-ng protein injection. Extracted ion chromatograms are shown for identified proteins along the elution gradient.

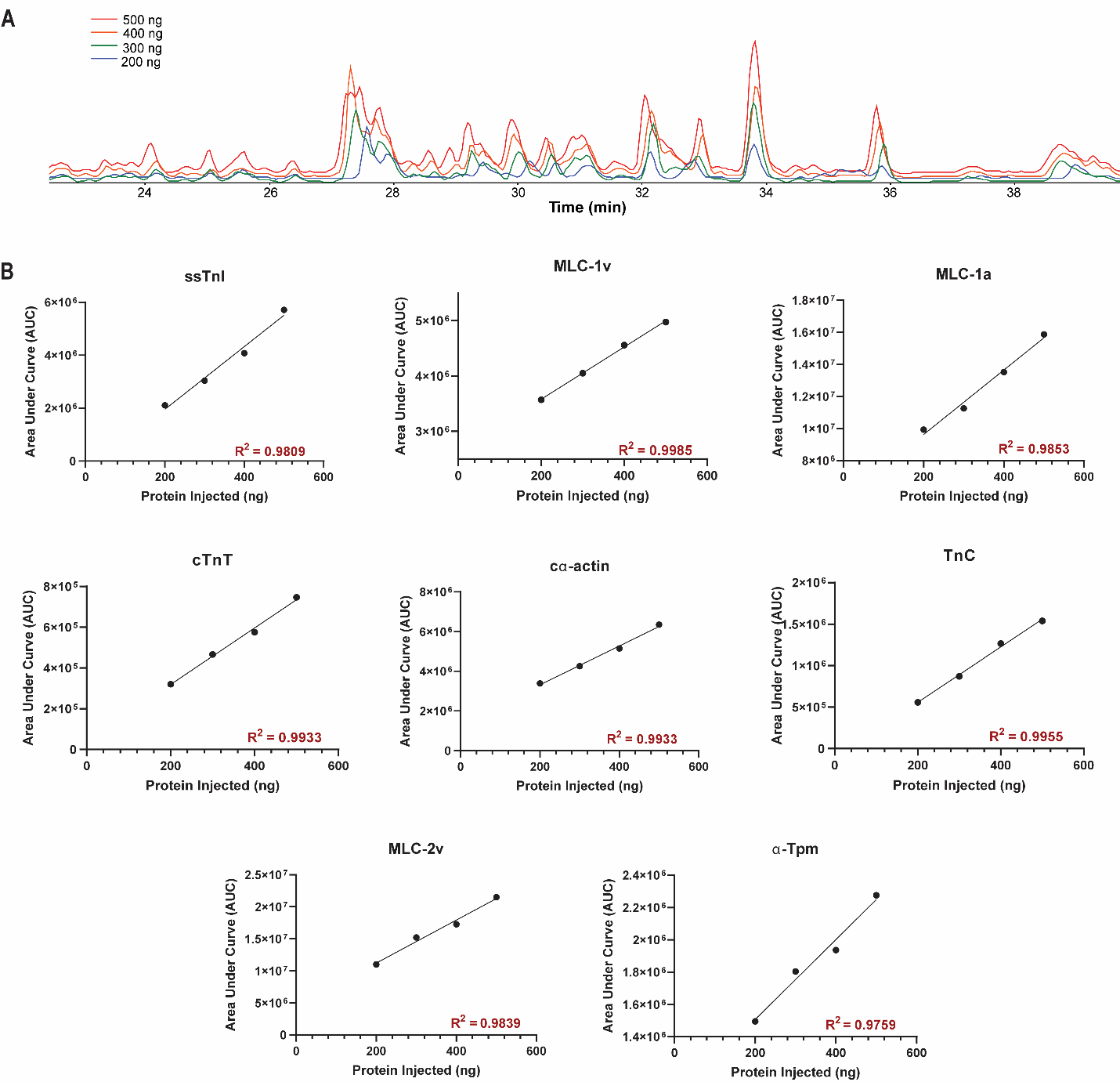

**Supplemental Figure 12. Instrument Linear Response for Intact Protein Analysis.** A) Various protein amounts injected on the IMPACT II to establish linear response curve – represented by base peak chromatogram. Trace color (total protein injected); Red (500 ng), Orange (400 ng), Green (300 ng), Blue (200 ng). B) Various linear curves generated for proteins identified. R^2^ values determined by simple linear regression.

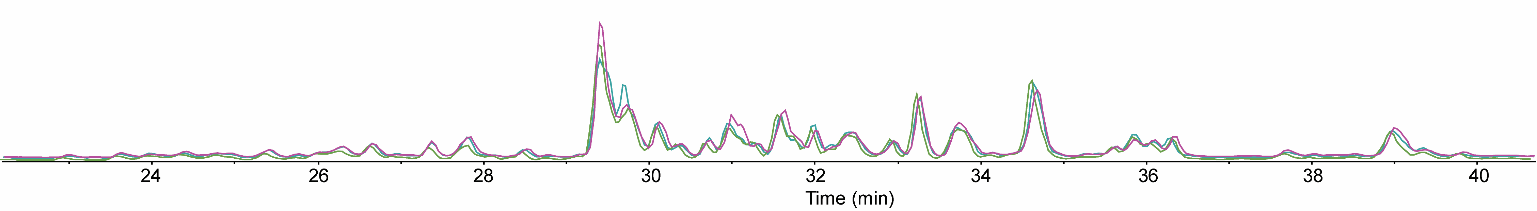

**Supplemental Figure 13. Instrument Stability using Base Peak Chromatograms.** Injections from example lactate injection shown from beginning (purple), middle (green), and end (blue) of instrument runs.

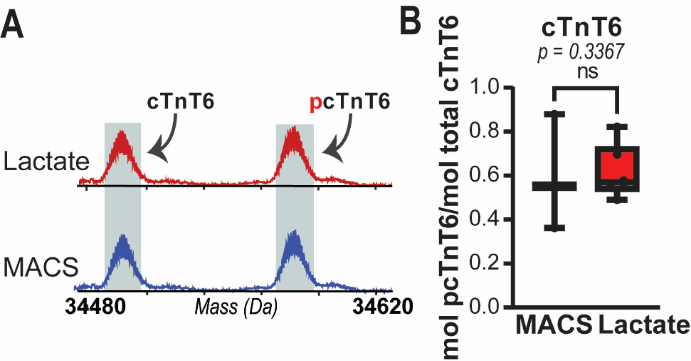

**Supplemental Figure 14. Phosphorylation of cTnT6. (**A) Deconvoluted mass spectra of cardiac troponin T 6 (cTnT) and its phosphorylated form. (B) pTotal calculations for cTnT6 (lactate = 0.616 ± 0.05 pTotal, n = 6 vs. MACS = 0.598 ± 0.15 pTotal, n = 3; unpaired t test, p = 0.8878). Unpaired t-test with alpha = 0.05.
